# Dizocilpine-derived K2060 exhibits enhanced blockage at NMDA receptors with impaired Mg^2+^ block and provides strong protection against audiogenic seizures

**DOI:** 10.64898/2026.10.06.756690

**Authors:** Marharyta Kolcheva, Jakub Netolicky, Anna Misiachna, Natalia Rybalko, Petra Zahumenska, Jovana Doderovic, Alena Pakhomova, Claudia Valentina De Sanctis, Gregory Klein Schneider, Katarina Hemelikova, Kristyna Rehakova, Stepan Kortus, Rostislav Turecek, Jan Korabecny, Ondrej Soukup, Marek Ladislav, Martin Horak

## Abstract

**Background and Purpose:** Voltage-dependent Mg^2+^ block is a defining feature of the GluN1/GluN2 subtype of *N*-methyl-D-aspartate receptors (NMDARs). We tested how impaired Mg^2+^ block at GluN1/GluN2A receptors mutated at the GluN2A-N615 position modifies inhibition by dizocilpine-derived K2060 compared with clinically used memantine and ketamine.

**Experimental Approach:** Electrophysiological recordings were performed on recombinant wild-type (WT) and mutated GluN1/GluN2A receptors expressed in HEK293T cells. Acute hippocampal slice electrophysiology and behavioural studies were conducted in WT and knock-in GluN2A-N615S mice. *In vivo* tolerability and efficacy were evaluated using open field testing, prepulse inhibition (PPI), and protection against audiogenic seizures (AGS).

**Key Results:** Mg^2+^ reduced the apparent potency of all three blockers and slowed onset at the GluN1/GluN2A-WT receptor. These Mg^2+^-dependent effects were absent or markedly reduced at the GluN1/GluN2A-N615S and GluN1/GluN2A-N615K receptors. In addition, K2060 showed slower unbinding and greater trapping than memantine. In hippocampal slices from GluN2A-N615S mice, K2060 exhibited enhanced but slower onset inhibition than memantine and ketamine. *In vivo*, K2060 caused no overt behavioural disruption at 5 mg/kg (*i.p.*) and did not further disrupt PPI. All three blockers protected GluN2A-N615S mice against AGS at 5 mg/kg (*i.p.*); only K2060 remained protective at 1 mg/kg (*i.p.*).

**Conclusions and Implications:** Impaired Mg²⁺ block removes a physiological constraint on the action of NMDAR blockers. K2060 exhibited the most favourable pharmacological profile, supporting its further evaluation for studies with NMDAR hyperfunction models. Additional pharmacokinetic, safety, and disease-model studies are warranted.

*What is already known?:*

- NMDAR blockers, including memantine, ketamine and the dizocilpine-derived K2060, bind within the ion channel pore, where Mg^2+^ can influence their potency and kinetics.
- The GluN2A-N615S variant reduces Mg^2+^ block and causes NMDAR hyperfunction associated with neuronal hyperexcitability and audiogenic seizure susceptibility in mice.

*What does this study add?:*

- Impaired Mg^2+^ block of the mutated NMDARs at the GluN2A-N615 position removes Mg^2+^-dependent modulation of blocker potency and kinetics.
- K2060 produces sustained inhibition of recombinant and endogenous GluN2A-N615S-containing NMDARs and protects GluN2A-N615S mice against AGS at a lower dose than memantine or ketamine.

*What is the clinical significance?:*

- *GRIN2A* pore variants that reduce Mg^2+^ block alter sensitivity to NMDAR blockers.
- K2060 warrants further preclinical evaluation for NMDAR hyperfunction animal models.

## 1 INTRODUCTION

The *N*-methyl-D-aspartate receptors (NMDARs) are ionotropic glutamate receptors with a central role in excitatory synaptic transmission in the mammalian central nervous system (CNS). NMDAR subunits are encoded by seven genes: *GRIN1* for GluN1, *GRIN2A–D* for GluN2A–D, and *GRIN3A–B* for GluN3A–B (Monyer et al., 1992; Paoletti et al., 2013; Hansen et al., 2021). In the adult forebrain, the major NMDAR populations contain GluN2A and/or GluN2B subunits and comprise diheteromeric GluN1/GluN2A and GluN1/GluN2B receptors (Figure 1), as well as triheteromeric GluN1/GluN2A/GluN2B receptors (Stroebel et al., 2018; Hansen et al., 2021; Zhang et al., 2025). The M2 re-entrant loops of the GluN1 and GluN2 subunits line the narrowest region of the ion channel and form its cation-selectivity filter. This region contains the conserved Q/R/N site, named for the glutamine, arginine, or asparagine residues occupying the homologous pore position across ionotropic glutamate receptor subfamilies (Hansen et al., 2021). Recent cryo-EM studies localized structural determinants of voltage-dependent Mg²⁺ block to this asparagine-lined selectivity-filter region (Huang et al., 2025), subsequently described as a narrow “Asn-cage” formed by GluN1-N616 and GluN2B-N615/N616, with GluN2A-N614/N615 occupying the homologous GluN2 positions (Huang et al., 2025; Steigerwald et al., 2026). Residues within and adjacent to this region critically determine ion selectivity and voltage-dependent Mg^2+^ block and contribute to the binding sites of clinically used open-channel blockers, including memantine and ketamine (Glasgow et al., 2018; Song et al., 2018; Chou et al., 2022; Zhang et al., 2021). It is well established that extracellular Mg^2+^ can decrease the potency of blockers and modify the kinetics of their inhibition at wild-type (WT) NMDARs (Sobolevsky et al., 1998; Kashiwagi et al., 2002; Kotermanski and Johnson, 2009; Glasgow et al., 2018; Kolcheva et al., 2021, 2023).

**FIGURE 1.**
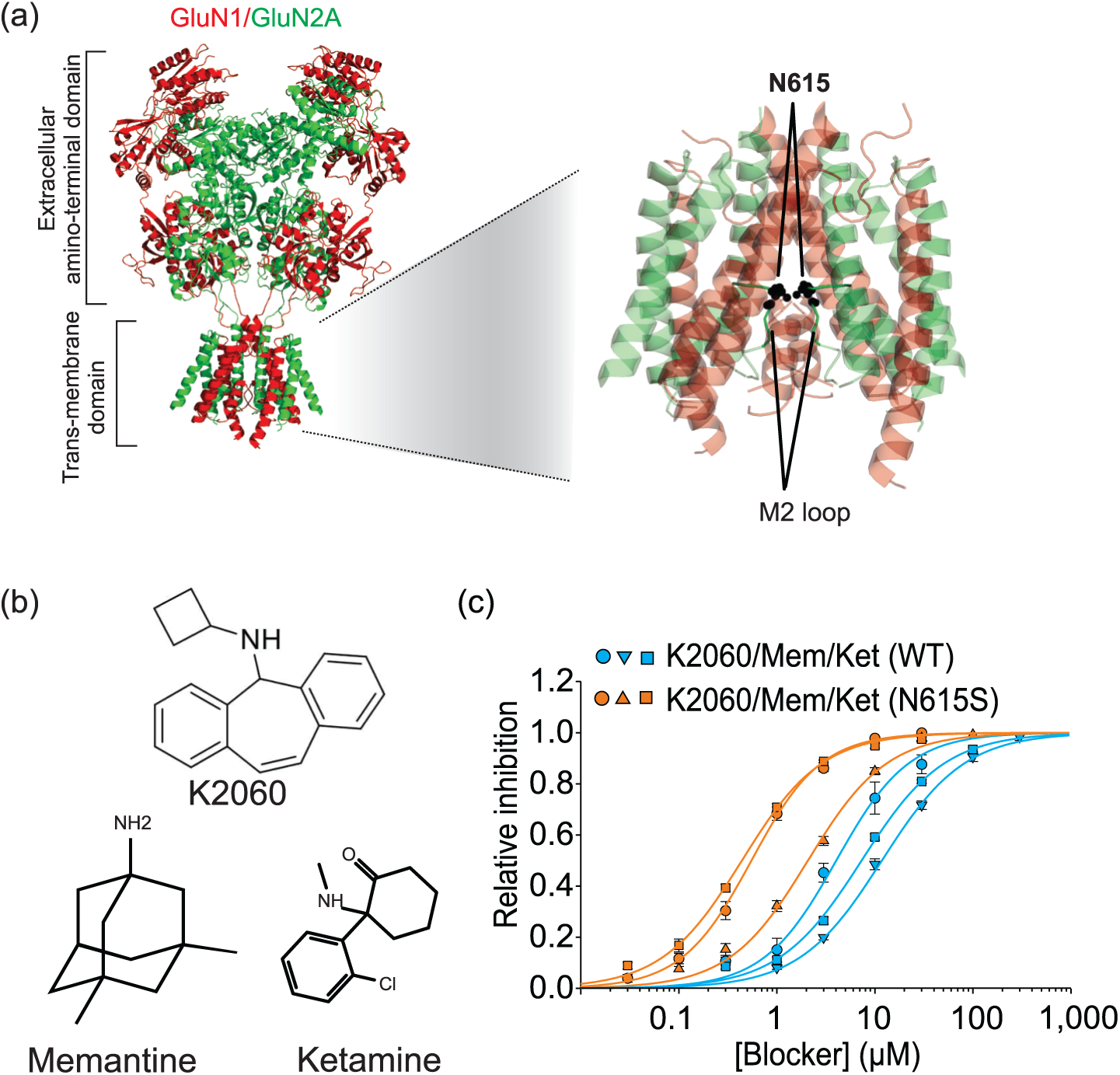
The N615S variant altered inhibition by K2060, memantine and ketamine in the presence of 1 mM Mg^2+^. (a) Structural view of the GluN1/GluN2A NMDAR (PDB: 7EOS), with GluN1 and GluN2A subunits shown in red and green, respectively. The full receptor is shown on the left, with the amino-terminal domain and transmembrane domain indicated. The enlarged transmembrane-domain view highlights the GluN2A-N615 residues within the ion channel pore-forming M2 loop. (b) Chemical structures of K2060, memantine and ketamine. (c) Concentration–response curves for inhibition of recombinant WT and N615S receptors by K2060, memantine and ketamine in the presence of 1 mM extracellular Mg^2+^. Concentration-response data are shown as mean ± SEM. IC_50_ values were determined by fitting concentration-response relationships from individual cell recordings using *Equation 1* and are reported in Table 1; full statistical details are provided in Supplementary Table S5.

**Table 1.**
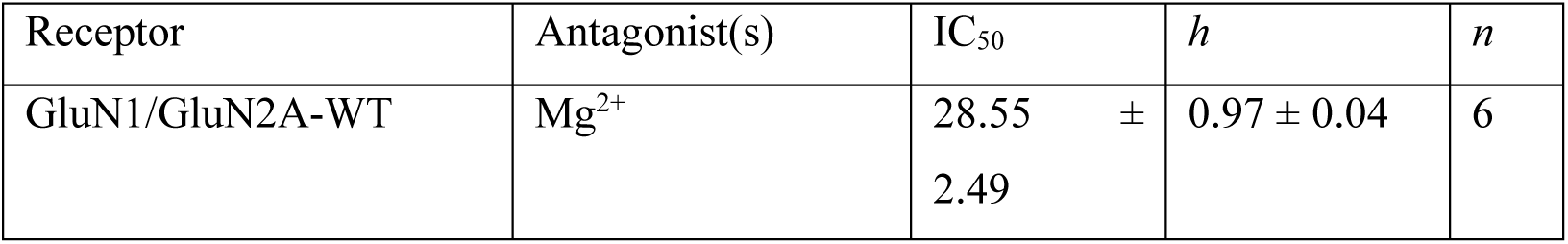
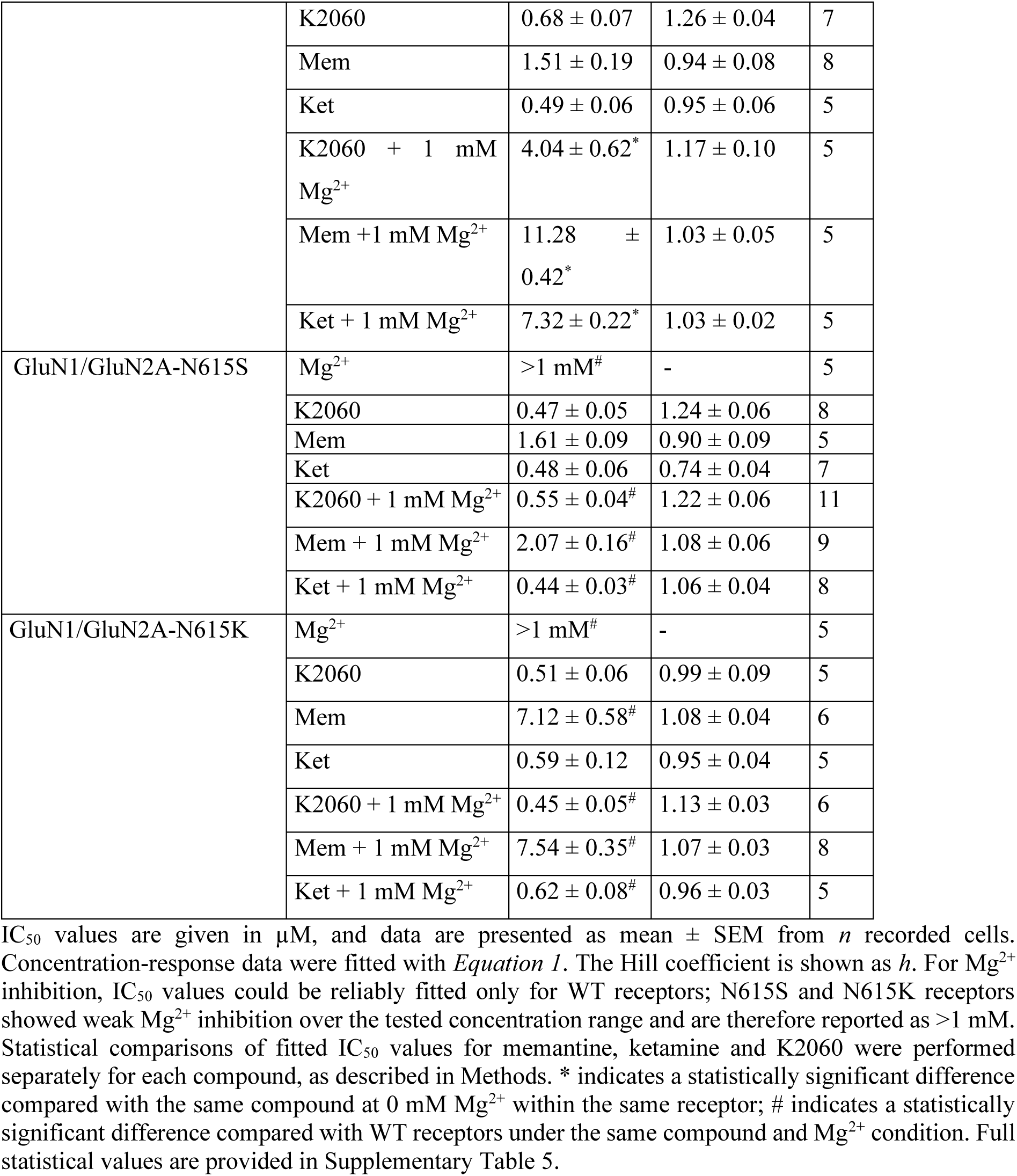
Fitting parameters for the concentration-response relationships for Mg^2+^, memantine, ketamine, and K2060 inhibition were measured for the indicated NMDARs expressed in HEK293T cells (related data are shown in Supplementary Figure S1a–n and Figure 6 a–b)

Consistent with the critical role of the M2-loop Q/R/N region and selectivity filter in ion permeation and voltage-dependent Mg²⁺ block, several disease-associated missense variants affecting this region have been reported in *GRIN1*, *GRIN2A*, and *GRIN2B* genes and are listed in the GRIN Variants Database (GRINdb; https://www.rarevariants.org/grindb/; accessed on 22 July, 2026). These include N616K, N616S, S617C, S617F, G618D, G618R, G618S, G618V, I619M, and G620R variants in *GRIN1* gene; N614S, N615K, S616F, and S616Y variants in *GRIN2A* gene; and N615I, N615K, N615S, N615Y, N616K, N616S, S617F, S617T, V618G, and V620M variants in *GRIN2B* gene. Among these variants, G620R in the *GRIN1* gene; N614S, N615K, and S616F in the *GRIN2A* gene; and N615I, N616K, N616S, V618G, and V620M in the *GRIN2B* gene have been reported in patients with epilepsy, seizures, or epilepsy-related disorders. Consistent with the distinct developmental expression profiles of GluN2 subunits, disease-associated variants in *GRIN2A* and *GRIN2B* genes are associated with partially overlapping but distinct spectra of CNS phenotypes (Paoletti et al., 2013; Platzer et al., 2017; Strehlow et al., 2019). For variants that impair voltage-dependent Mg²⁺ block, excessive NMDAR activation at hyperpolarised membrane potentials may promote excessive NMDAR-mediated excitation and contribute to neuronal hyperexcitability (Fedele et al., 2018). However, variants in this region may also alter NMDAR expression, ion permeability, current amplitude, agonist sensitivity, and channel kinetics; therefore, impaired Mg²⁺ block should not be considered the sole pathogenic mechanism without variant-specific functional evidence (Fedele et al., 2018; Korinek et al., 2024). NMDAR blockers may consequently represent a therapeutic strategy for selected variants with an overall gain-of-function phenotype, although their efficacy is likely to depend on both the specific variant and the pharmacological properties of the blocker (Platzer et al., 2017; Fedele et al., 2018; Strehlow et al., 2019; Benke et al., 2021). Progress in this area remains limited because only a small proportion of patient-associated variants have been investigated in neuronal preparations or animal models, and systematic comparative *in vitro* and *in vivo* studies of multiple NMDAR blockers remain scarce (Benke et al., 2021; Korinek et al., 2024).

Here, we employed a gene-targeted mouse line carrying the GluN2A-N615S (N615S) variant, located within the Q/R/N region at the tip of the M2 loop (Bertocchi et al., 2021). N615S mice provide an experimentally tractable model of impaired voltage-dependent Mg²⁺ block and exhibit pronounced exploratory hyperactivity, altered attention-dependent learning, and high susceptibility to audiogenic seizures (AGS) associated with hyperexcitability in midbrain circuits (Bertocchi et al., 2021, 2024). We characterised the *in vitro* and *in vivo* effects of K2060, a dizocilpine-derived potent and reversible NMDAR blocker (Misiachna et al., 2024), and directly benchmarked them against those of the clinically established blockers memantine and ketamine. Importantly, we showed that K2060 protects N615S mice against AGS at a lower concentration than memantine and ketamine, and that the blocker effects observed for GluN1/GluN2A-N615S receptors extend to the GluN1/GluN2A receptors carrying the disease-associated GluN2A-N615K variant.

## 2 METHODS

### 2.1 Animals

All experiments involving animals or animal tissue were performed in accordance with national and institutional regulations for the care and use of laboratory animals and were approved by the Ethical Committee of the Institute of Experimental Medicine, CAS. Animal experiments were designed and reported in accordance with ARRIVE 2.0 guidelines (Percie du Sert et al., 2020) and the recommendations of the *British Journal of Pharmacology* for animal experimentation and data analysis (Lilley et al., 2020). Experiments were performed using wild-type C57BL/6N mice (WT mice) or homozygous B6.129-*Grin2A*tm2Rsp/Kctt mice carrying the GluN2A-N615S variant (N615S mice) (RRID:IMSR_EM:09319, Bertocchi et al., 2021) purchased from the EMMA European Mutant Mice Archive (EM:09319:B6.129-Grin2atm2Rsp/Kctt; https://www.informatics.jax.org/allele/MGI:6406404). All animals were bred and housed under standard conditions (12 h light/dark cycle, controlled temperature and humidity, food and water *ad libitum*), in groups of two to four per cage.

For acute hippocampal slice recordings, mice of both sexes were used at postnatal days 35–42 (P35–P42). For behavioural experiments, male mice were used at 2–3.5 months of age. Behavioural experiments were performed in male mice only because the present study was designed as an initial *in vivo* pharmacological comparison and was not powered to include sex as an additional biological variable. Previous studies using N615S mice included both sexes and showed that the main behavioural phenotype, including susceptibility to audiogenic seizures, is present in both male and female mice (Bertocchi et al., 2021, 2024). Although sex may influence the distribution of WR, AGS, and AGS-RA stages, these differences appear less relevant to the dichotomous normal-versus-abnormal endpoint used in our AGS analysis. A formal sex-stratified pharmacological analysis would require larger cohorts and was not pursued in line with the reduction of animal use. The male-only design, therefore, represents a limitation of the behavioural component of the study.

### 2.2 Molecular biology and HEK293T cell culture

Human cDNA constructs encoding GluN1-4a (GluN1) and GluN2A subunits were used as described previously (Kolcheva et al., 2021). Point mutations in the coding region of the GluN2A subunit were introduced using the QuikChange site-directed mutagenesis kit (Agilent Technologies, Santa Clara, CA, USA), and all constructs were verified by DNA sequencing. HEK293T (RRID:CVCL_0063) cells were maintained in Opti-MEM I medium supplemented with 5% fetal bovine serum (FBS; Thermo Fisher Scientific, Waltham, MA, USA). For transient expression, cells grown in 24-well plates were transfected in 50 μl Opti-MEM I using 0.9 μl PolyMag transfection reagent (OZ Biosciences, Marseille, France). Each well received a total of 0.9 μg of cDNA encoding the GluN1 and GluN2A subunits, together with GFP (pQBI 25; Takara Bio Inc., Kusatsu, Japan), to identify transfected cells. After transfection, cells were trypsinised and replated at low density onto poly-L-lysine-coated glass coverslips of 3 cm diameter. To reduce excitotoxicity caused by NMDAR expression, cells were maintained in Opti-MEM I supplemented with 1% (v/v) FBS, 2 mM MgCl_2_, 1 mM DL-2-amino-5-phosphonovaleric acid and 3 mM kynurenic acid until electrophysiological recordings.

### 2.3 Electrophysiology

#### 2.3.1 Electrophysiology in HEK293T cells

Whole-cell patch-clamp recordings were performed from the transfected HEK293T cells 16–48 h after the transfection using an Axopatch 200B amplifier (Axon Instruments, Inc., Foster City, CA) with series resistance (<10 MΩ) and ∼80% capacitance compensation at a holding membrane potential of −60 mV. The current responses were low-pass filtered at 2 kHz using an eight-pole Bessel filter, digitised at 5 kHz, and recorded using pCLAMP 11 (Molecular Devices, San Jose, CA, USA; RRID:SCR_011323). Borosilicate glass pipettes with a tip resistance of 3–6 MΩ were prepared using a P-1000 puller (Sutter Instrument, Novato, CA, USA) and filled with intracellular solution (ICS) containing (in mM): 125 gluconic acid, 15 CsCl, 5 EGTA, 10 HEPES, 3 MgCl_2_, 0.5 CaCl_2_, and 2 ATP-Mg salt (pH 7.2 with CsOH). The extracellular recording solution (ECS) contained (in mM) 160 NaCl, 2.5 KCl, 10 HEPES, 10 D-glucose, 0.01 EDTA, and 1 CaCl_2_ (pH 7.3 with NaOH). The ECS contained a saturating concentration of 50 µM glycine, which is not indicated in the figures. The NMDAR current responses were evoked by a saturating concentration of 1 mM L-glutamate in the ECS, using a multi-barrel rapid application system with a solution exchange time constant of ∼20 ms at the measured cell (Kolcheva et al., 2021).

The concentration-response curves for open-channel blockers were obtained using the following equation:

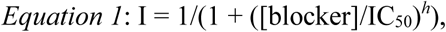

IC_50_ is the concentration of memantine, ketamine, or K2060 that produced 50% inhibition of the agonist-evoked current, [blocker] is the concentration of memantine, ketamine, or K2060, and *h* is the apparent

Hill coefficient. The time course of blocker inhibition was analysed using a double exponential function, and weighted time constants (τ_w_) were obtained using the following equation:

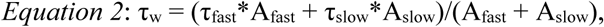

the faster component has a time constant τ_fast_ and an amplitude A_fast_, and the slower component has a time constant τ_slow_ and amplitude A_slow_; normalised values of A_fast_ were obtained by dividing each A_fast_ value by (A_fast_ + A_slow_) (Glasgow et al., 2018).

To assess the degree of blocker trapping, we employed a well-established double-pulse protocol (Bolshakov et al., 2003; Kotermanski et al., 2009; Bettini et al., 2022). NMDARs were activated with 30 μM NMDA and 10 μM glycine, and after steady-state current response was achieved, a saturating concentration (100 μM) of K2060, memantine or ketamine was applied to induce a complete block. This was followed by a wash with ECS for a period corresponding to 10-fold the τ_off_ values determined for agonist-present unbinding; NMDARs were then reactivated by co-application of 30 μM NMDA and 10 μM glycine. The recovery traces were fit with a double-exponential function, thereby separating the fast and slow unbinding components. The fast component was interpreted as the fraction of blocker released during the wash period with ECS, while the slow component reflected the fraction of trapped blocker. L-glutamate was avoided as an agonist because its rapid offset prevents reliable separation of the blockers’ fast and slow unbinding phases.

#### 2.3.2 Electrophysiology in acute hippocampal slices

Mouse hippocampal slices were prepared from young adult WT or N615S mice of both sexes (postnatal day 35–42). Animals were deeply anaesthetised with inhaled isoflurane, decapitated, and their brains rapidly removed and immersed in ice-cold, oxygenated artificial cerebrospinal fluid (aCSF; 95% O_2_/5% CO_2_). The aCSF contained (in mM): 124 NaCl, 3.5 KCl, 1.25 NaH_2_PO_4_, 26 NaHCO_3_, 4 MgSO_4_, 0.5 CaCl_2_, and 11 D-glucose (pH 7.3). Horizontal 300-μm slices were cut in ice-cold aCSF, hemisected, and incubated at 32 °C for 30 min in oxygenated aCSF containing 1 mM MgSO_4_ and 1 mM CaCl_2_ (300– 310 mOsm/kg). Slices were subsequently equilibrated to room temperature for at least 60 min before recordings.

Whole-cell voltage-clamp recordings were performed from CA1 pyramidal neurons using a Multiclamp 700B amplifier (Molecular Devices, San Jose, CA, USA). Signals were sampled at 25 kHz (Digidata 1550B; Molecular Devices, San Jose, CA, USA) and low-pass filtered at 10 kHz using pCLAMP 11. CA1 pyramidal neurons were visualised using Dodt gradient contrast with a 40× water-immersion objective, an Axiocam 305 colour digital camera and an Axio Examiner.Z1 microscope (Carl Zeiss Microscopy GmbH, Jena, Germany). Recordings were performed at 32 ± 1 °C in oxygenated aCSF. Each slice was randomly selected, and only one cell was recorded per slice. Patch pipettes (4–5 MΩ) were pulled from thin-walled borosilicate glass and filled with ICS (in mM): 120 Cs-gluconate, 9 NaCl, 10 HEPES, 10 BAPTA, 4 Mg-ATP, 0.3 Mg-GTP, and 2 MgCl_2_ (pH adjusted to 7.3 with CsOH; osmolarity ∼290 mOsm/kg). After a 10-min equilibration period to allow ICS diffusion, evoked NMDAR-mediated excitatory postsynaptic currents (NMDAR-eEPSCs) were recorded at a holding potential of −60 mV (liquid junction potential was not corrected). To isolate NMDAR-eEPSCs, 10 µM NBQX (AMPAR antagonist) and 100 µM picrotoxin (GABA_A_ receptor antagonist) were continuously present. Bipolar stimulus (100 µs/phase biphasic pulse) at 0.05 Hz was applied via a concentric bipolar tungsten electrode (#TM33CCINS-B; World Precision Instruments, Sarasota, FL, USA) placed in the stratum radiatum using an STG5 stimulator (Multi Channel Systems, Reutlingen, Germany) triggered by Clampex 11. Each stimulus contained a train of five stimuli at 50 Hz.

The onset of blocker-induced inhibition was assessed using eEPSCs. Each experiment was preceded by at least 7 min of stable baseline stimulation to ensure response stability, and blocker onset was recorded for up to 55 min. The current amplitude evoked by the fifth stimulus in each train was analysed in Clampfit 11. For visualisation, three stimulation trains per min were averaged, resulting in one data point per min. To evaluate the onset of inhibition, current amplitudes were first expressed as a relative response to baseline. Specifically, for each time point during blocker application, the evoked current (I_t_) was normalised (I_norm(t)_) to the mean current during the 6-min baseline preceding blocker onset (I_baseline_), producing a trace of relative inhibition over time. This representation was used to depict the inhibition profile as a fraction of the pre-blocker baseline.

To further evaluate the kinetics of blocker-induced inhibition, the baseline-normalised trace was transformed to reflect the progression toward maximal inhibition. The mean value of the final 5 min of blocker application (I_min_) was taken as the asymptotic maximum inhibition. The normalised inhibition kinetics were then computed using the following transformation:

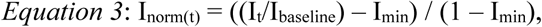

where I_t_ is the evoked current amplitude at time point t, I_baseline_ is the mean current amplitude during the 6-min pre-blocker baseline, I_min_ is the mean of the last 5 min of blocker, and I_norm(t)_ represents the fraction of inhibition achieved at time t, scaled such that 1 corresponds to baseline (no inhibition) and 0 to maximal inhibition.

This two-step approach allowed visualisation of raw inhibition and the kinetic progression toward steady-state inhibition. I_norm(t)_ traces were subsequently fitted with a single-exponential decay to extract kinetic time constants. To examine potential presynaptic effects, peak current amplitudes from the first and second stimuli were also analysed and compared with those of the fifth stimulus. Series resistance and cell capacitance were continuously monitored throughout recordings. Cells with >20% variation in these parameters were excluded from analysis. Series resistance compensation was not applied.

### 2.4 Acoustic startle response (ASR) and prepulse inhibition (PPI)

For ASR and PPI testing, male WT and N615S mice (2–3.5 months old) were placed in a ventilated sound-attenuated chamber (Habitest, model E10-21; Coulbourn Instruments, Holliston, MA, USA). During the test, animals were confined in a small wire cage positioned on a motion-sensitive platform.

Whole-body startle responses were detected with a load-cell transducer (E45-11; Coulbourn Instruments, Holliston, MA, USA), amplified, and processed using a System III RP2.1 processor (Tucker-Davis Technologies, Alachua, FL, USA). A continuous 70 dB SPL background noise was present throughout the experiments, unless otherwise indicated. To minimise stress, cage bedding was not changed on the day before or during testing, and mice were gently transferred to the chamber immediately before the experiment.

PPI was assessed using trials consisting of a startle pulse alone, a prepulse alone, a prepulse–pulse combination or no stimulus. Startle pulses were delivered at 120 dB SPL. Prepulses were delivered at 74, 78, 82 or 86 dB SPL for 50 ms and, in prepulse–pulse trials, preceded the startle pulse by 50 ms. No-stimulus trials consisted of background noise only. Each session began with a habituation period in background noise, followed by a short series of isolated startle pulses. The full PPI session then consisted of 10 distinct trial types, including pulse-alone, prepulse-alone, prepulse–pulse combinations at four prepulse intensities, and no-stimulus trials. Each trial type was presented 10 times in pseudorandom order, resulting in 100 trials per session, with inter-trial intervals of 15–30 s.

Testing was performed over three consecutive days. On day 1, mice were acclimatised to the testing chamber and acoustic stimulation protocol. On day 2, the PPI protocol was performed to obtain baseline values for each animal. On day 3, the same PPI protocol was repeated 30 min after intraperitoneal (*i.p.*) administration of K2060 or vehicle. K2060 was freshly prepared in saline containing 5% DMSO and 5% Kolliphor, and the corresponding vehicle was the same solution without K2060. K2060 was administered at 5 mg/kg.

Startle responses were quantified within a 100-ms window after pulse onset. The extent of PPI was expressed as:

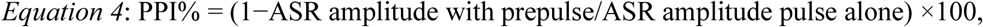

PPI% = (1−ASR amplitude with prepulse/ASR amplitude pulse alone) ×100, where higher values indicate stronger prepulse-mediated inhibition of the startle response. This paradigm allowed us to evaluate potential sensorimotor gating deficits induced by the selected dose of K2060.

### 2.5 Open field test (OFT)

Male WT and N615S mice aged 2–3.5 months were used for OFT experiments. K2060 was freshly prepared in saline containing 5% DMSO and 5% Kolliphor, and the corresponding vehicle was the same solution without the blocker. Mice were injected with vehicle or K2060 at doses of 5, 15, 30, or 50 mg/kg in a 10 ml/kg volume (*i.p.*) and were immediately placed individually into a 50 × 50 cm open-field arena for a 40-min trial. Behavioural data were collected and processed using EthoVision XT (RRID: SCR_000441; Noldus, Wageningen, the Netherlands). The total distance travelled in the arena was evaluated as a measure of hyperlocomotion (Seibenhener and Wooten, 2015). In addition, video recordings were inspected for overt acute behavioural signs typically associated with NMDAR antagonists, including stereotypic or repetitive behaviours, abnormal posture, and impaired motor coordination. Between trials, the arena was cleaned with 70% ethanol to remove residual odours.

### 2.6 Audiogenic seizure (AGS) induction protocol

The AGS was induced in male N615S mice (2–3.5 months old) using a protocol adapted from a previous study (Bertocchi et al., 2021). Mice were placed individually into a custom-built, soundproof chamber. Acoustic stimulation (11 kHz, 110 dB SPL) was generated using a white noise generator (RFT 03004; Robotron, Germany), processed through RFT 01013 filters, amplified with a custom-made amplifier, and delivered via a DE700 speaker (B&C Speakers; Florence, Italy) mounted on a horn. Sound pressure levels were calibrated using a B&K 4939 1/4 microphone and a B&K 2231 sound level meter (Hottinger Brüel & Kjaer, Naerum, Denmark). The stimulus was presented for up to 80 sec, in four 20-s bouts separated by 2-s breaks, or until the onset of AGS. Mice were habituated in the chamber for 5 min before stimulation. Behavioural responses were recorded using a webcam. If seizures did not occur or if death did not follow the seizure episode, mice remained undisturbed in the chamber for an additional 5 min after tone presentation. Blockers were freshly prepared in saline containing 5% DMSO and 5% Kolliphor. Mice received injections of 1 or 5 mg/kg of memantine, ketamine, K2060, or saline (*i.p.*) 30 min before acoustic stimulation. Behavioural classification during sound exposure was performed according to previously described criteria (Bertocchi et al., 2024). The typical response consisted of wild running (WR), followed by loss of posture with clonic and tonic limb extension (AGS), and in some cases, respiratory arrest and death (AGS-RA), occurring in this specific sequence.

### 2.7 Data and statistical analysis

Experimental design, data analysis, and statistical procedures were performed in accordance with the recommendations of the *British Journal of Pharmacology* (Curtis et al., 2022). Data are presented as mean ± SEM unless stated otherwise, and individual observations are shown where possible. The experimental unit was defined before analysis. For HEK293T cell experiments, *n* represents the number of recorded cells. For hippocampal slice experiments, *n* represents cells/slices, with one neuron recorded per slice. For behavioural experiments, *n* represents the number of animals.

Group sizes were chosen based on previous studies using the N615S mice and related NMDAR pharmacology experiments, while applying the principle of reducing animal use. No formal a priori power calculation was performed. Electrophysiological experiments were designed to obtain at least five independent recordings per group whenever technically feasible. In a small number of technically demanding, long-duration recovery or trapping experiments, final group sizes of n = 4 were examined after predefined quality-control exclusions. These datasets are shown with individual data points and exact *n* values and were interpreted cautiously in the context of the overall pharmacological pattern.

Animals were allocated to treatment groups using a randomised design, and the order of blocker administration was balanced across groups where possible. Behavioural scoring of AGS responses was performed from video recordings according to predefined criteria. Experimenters performing behavioural scoring and data analysis were blinded to treatment group.

Inclusion and exclusion criteria were defined before analysis. Electrophysiological recordings were excluded if series resistance or cell capacitance changed by ≥20% during the experiment or if a predefined technical failure occurred. Behavioural data were excluded only in cases of technical failure, such as unsuccessful injection, equipment malfunction or incomplete video recording. No animals or data points were excluded based on biological response alone.

Statistical analyses were performed using SigmaStat 4.0 (Systat Software Inc., San Jose, CA, USA). Normality was assessed using the Shapiro–Wilk test, and homogeneity of variance was assessed prior to parametric analyses. For two-group comparisons, Student’s t-test, Welch’s t-test, Mann–Whitney rank-sum test or Wilcoxon signed-rank test was used as appropriate. For comparisons involving more than two groups or factors, one-way ANOVA, two-way ANOVA or two-way repeated-measures ANOVA was used, followed by Holm–Sidak *post hoc* tests where appropriate. When assumptions for parametric testing were not fully met in repeated-measures analyses, ANOVA results were reported as descriptive summaries and interpreted with caution. Exact tests used for individual datasets are provided in the corresponding supplementary statistical tables.

For concentration–response analyses, IC_50_ values were obtained by fitting individual concentration–response relationships from single recorded cells. Statistical comparisons of fitted IC_50_ values were performed separately for each blocker using two-way ANOVA when assumptions were met or predefined pairwise Welch’s t-tests or Mann–Whitney rank-sum tests with Holm correction when assumptions were not met. Time-course inhibition experiments in acute hippocampal slices were analysed separately for each blocker and normalisation approach. Baseline-normalised responses were used to compare the overall development of inhibition, whereas scaled responses were used to compare the apparent onset profile independently of final inhibition magnitude. Final 5-min inhibition was analysed by two-way ANOVA with genotype and blocker as factors.

Categorical behavioural outcomes in the AGS experiments were analysed using Fisher’s exact test. For statistical analysis, outcomes were dichotomised as normal response versus abnormal audiogenic response. Wild running (WR), audiogenic seizure (AGS), and AGS followed by respiratory arrest (AGS-RA) were pooled into the abnormal response category. Each treatment group was compared with the corresponding dose-matched vehicle group. Differences were considered statistically significant when P < 0.05.

### 2.8 Materials

Reagents used for preparing ECS, ICS, aCSF, glutamate, glycine, NMDA, DL-2-amino-5-phosphonovaleric acid, kynurenic acid, picrotoxin and NBQX were obtained from Merck (Darmstadt, Germany), Tocris Bioscience (Bristol, UK) or Hello Bio (Bristol, UK). Cell-culture media and FBS were from Thermo Fisher Scientific (Waltham, MA, USA). PolyMag transfection reagent was from OZ

Biosciences (Marseille, France). The QuikChange site-directed mutagenesis kit was from Agilent Technologies (Santa Clara, CA, USA), and pQBI 25 plasmid containing GFP was from Takara Bio Inc. (Kusatsu, Japan). Memantine and ketamine were obtained from Hello Bio (Bristol, UK), and K2060 was synthesised as described (Misiachna et al., 2024).

### 2.9 Nomenclature of targets and ligands

Key protein targets and ligands are hyperlinked to corresponding entries in http://www.guidetopharmacology.org, the common portal for data from the IUPHAR/BPS Guide to PHARMACOLOGY (Harding et al., 2026) and are permanently archived in the Concise Guide to PHARMACOLOGY 2025/26 (Alexander et al., 2025).

## 3 RESULTS

### 3.1 The GluN2A-N615S variant reduced Mg^2+^-dependent reduction in apparent potency of blockers

First, we asked whether the GluN2A-N615S variant (N615S variant) alters the apparent inhibitory potency of K2060, memantine and ketamine, particularly under experimental conditions in which extracellular Mg^2+^ limits blocker access to wild-type GluN1/GluN2A (WT) receptors (Figure 1a,b). We therefore examined blocker inhibition at recombinant WT and GluN1/GluN2A-N615S (N615S) receptors expressed in the HEK293T cells, in the absence (0 Mg^2+^) and presence of 1 mM Mg^2+^. Consistent with previous data (Wollmuth et al., 1998; Bertocchi et al., 2021), N615S receptors showed reduced Mg^2+^ block (Table 1; Figure S1a,b). In 0 Mg^2+^, the inhibitory potencies of K2060, memantine, and ketamine were comparable between WT and N615S receptors. In contrast, in 1 mM Mg^2+^, the WT receptors showed the expected rightward IC_50_ shift for all three blockers, whereas the N615S receptors did not, thereby eliminating the Mg^2+^-dependent loss of apparent potency for all three blockers (Table 1; Figure 1c; Figure S1c-n).

### 3.2 The GluN2A-N615S variant altered inhibitory kinetics of blockers at the GluN1/GluN2A receptors in the presence of Mg^2*+*^

Because the inhibitory efficacy of a blocker is determined not only by its potency, we next assessed whether the N615S variant alters inhibition onset (τ_on_) and recovery (τ_off_) kinetics of K2060, memantine, and ketamine at 10, 30, and 100 μM in the absence and presence of Mg^2+^. As expected, increasing concentration accelerated τ_on_ (10 > 30 > 100 μM) at both WT and N615S receptors in 0 and 1 mM Mg^2+^. For K2060, τ_on_ was ∼3-fold slower at N615S than WT receptors in 0 Mg^2+^. Mg^2+^ further delayed τ_on_ at WT receptors but had no effect at N615S receptors, indicating that the Mg^2+^-dependent delay is abolished by the N615S variant (Figure 2b; Figure S1a; Table S1). By comparison, in 0 Mg^2+^, τ_on_ for memantine and ketamine was similar between WT and N615S receptors. In 1 mM Mg^2+^, τ_on_ slowed at WT receptors for memantine and ketamine, consistent with a Mg^2+^-dependent delay, whereas N615S receptors showed no Mg²⁺-dependent effect (Figure 2e, h; Figure S2b,c; Tables S2–3).

**FIGURE 2.**
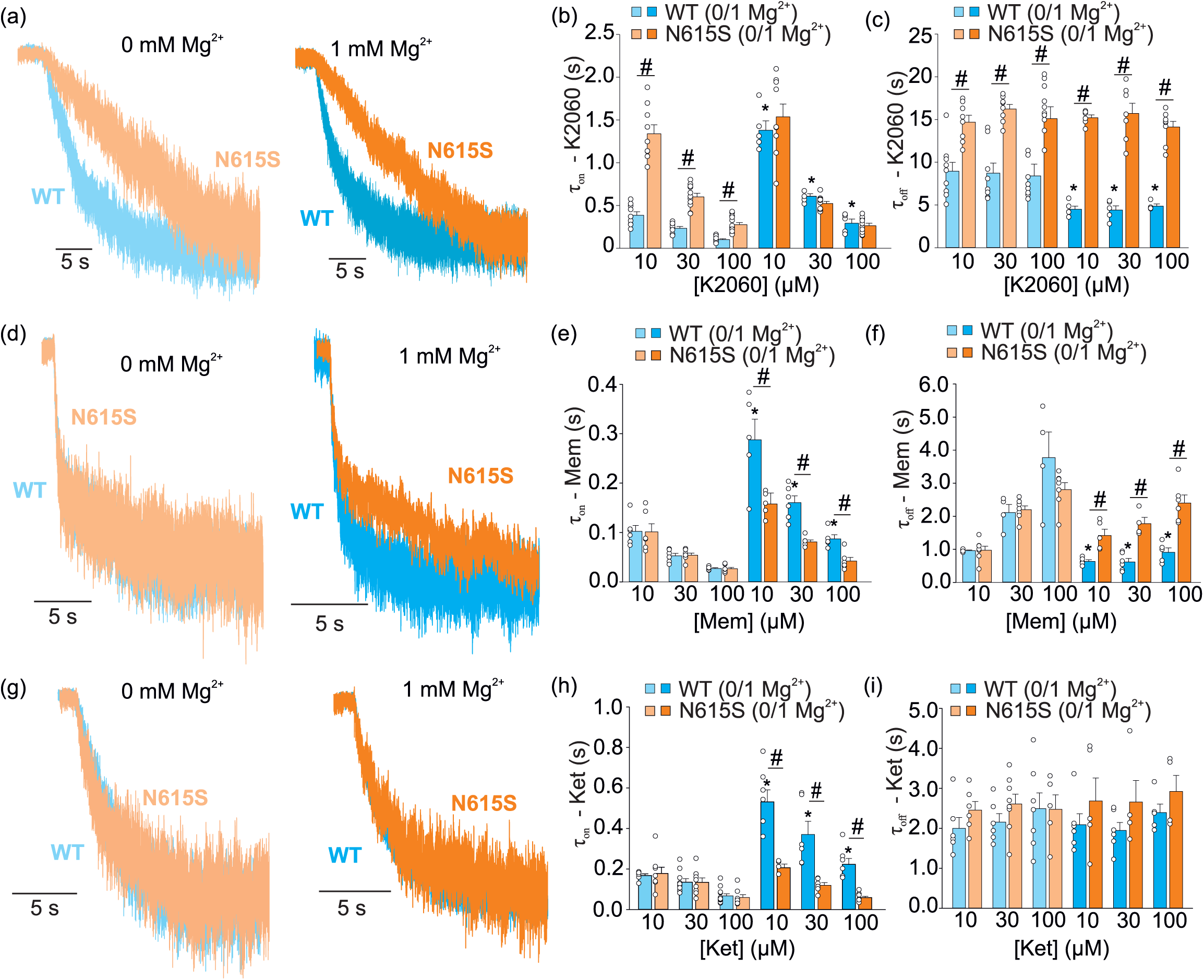
The N615S variant altered Mg^2+^-dependent inhibition kinetics of K2060, memantine and ketamine. (a, d, g) Representative current traces showing recovery from inhibition after removal of 30 μM K2060 (a), memantine (d) or ketamine (g) from recombinant WT and N615S receptors. Traces were recorded in 0 or 1 mM extracellular Mg^2+^, as indicated above each pair of traces. Blue traces show WT receptors and orange traces show N615S receptors; lighter colours indicate 0 mM Mg^2+^ and more saturated colours indicate 1 mM Mg^2+^. Traces were aligned to the time of blocker removal to compare recovery kinetics. (b, e, h) Summary of inhibition onset time constants (τ_on_) for K2060 (b), memantine (e) and ketamine (h) at 10, 30 and 100 μM in 0 and 1 mM extracellular Mg^2+^. (c, f, i) Summary of recovery time constants after blocker washout (τ_off_) for K2060 (c), memantine (f) and ketamine (i) at 10, 30 and 100 μM in 0 and 1 mM extracellular Mg^2+^. Kinetic time constants were obtained from exponential fits, and weighted recovery time constants were calculated using *Equation 2*. Data are presented as mean ± SEM, with individual recorded cells shown as open circles. * indicates a statistically significant difference compared with the same receptor and compound concentration in 0 mM Mg^2+^; # indicates a statistically significant difference between WT and N615S receptors under the same compound concentration and Mg^2+^ condition. Full kinetic values and *n* are provided in Supplementary Tables S1– S3. Statistical comparisons were performed as described in Methods, and full statistical values are provided in Supplementary Table S6.

Concerning τ_off_ kinetics, K2060 was concentration-independent but markedly slower than memantine and ketamine at WT receptors in 0 Mg^2+^ (Figure 2a,c; Figure S2a; Table S1). Memantine showed a concentration-dependent prolongation, whereas ketamine was concentration-independent, consistent with prior work (Glasgow et al., 2018). The same concentration pattern was evident at N615S receptors in 0 Mg^2+^. In 1 mM Mg^2+^, τ_off_ at WT receptors accelerated for memantine and K2060 relative to 0 Mg^2+^ and became concentration-insensitive, while τ_off_ for ketamine remained unchanged. At N615S receptors, τ_off_ was Mg^2+^-insensitive and preserved the concentration pattern observed in 0 Mg^2+^. In 0 Mg^2+^, τ_off_ for K2060 was markedly slower at N615S than WT receptors, whereas τ_off_ for memantine and ketamine were comparable. In 1 mM Mg^2+^, τ_off_ for K2060 and memantine were prolonged, while τ_off_ for ketamine was similar between WT and N615S receptors (Figure 2a–i; Figure S2a-c; Tables S1–3). Together, our data showed that K2060 exhibits the slowest recovery kinetics and the most prominent variant-dependent differences among tested blockers.

3.3 Trapping of K2060 and comparator blockers was not altered by GluN2A-N615S variant or Mg^2+^

Because the degree of trapping is also relevant to how blockers act at NMDARs, we next used a previously established double-pulse protocol for K2060, memantine and ketamine. All three blockers have been described as partial (or nearly full) trappers (Blanpied et al., 1997; Mealing et al., 1999; Kotermanski et al., 2009; Misiachna et al., 2024). We found no differences in the degree of trapping of all three blockers between WT and N615S receptors, either in 0 Mg^2+^ or 1 mM Mg^2+^ (Figure 3a–h; Table S4). At WT receptors, K2060 and ketamine showed greater trapping than memantine. The same rank order was held at N615S receptors, with memantine showing consistently lower trapping than the other two blockers. Thus, K2060 and ketamine are more likely than memantine to remain trapped in the ion channel pore during repeated cycles of NMDAR activation.

**FIGURE 3.**
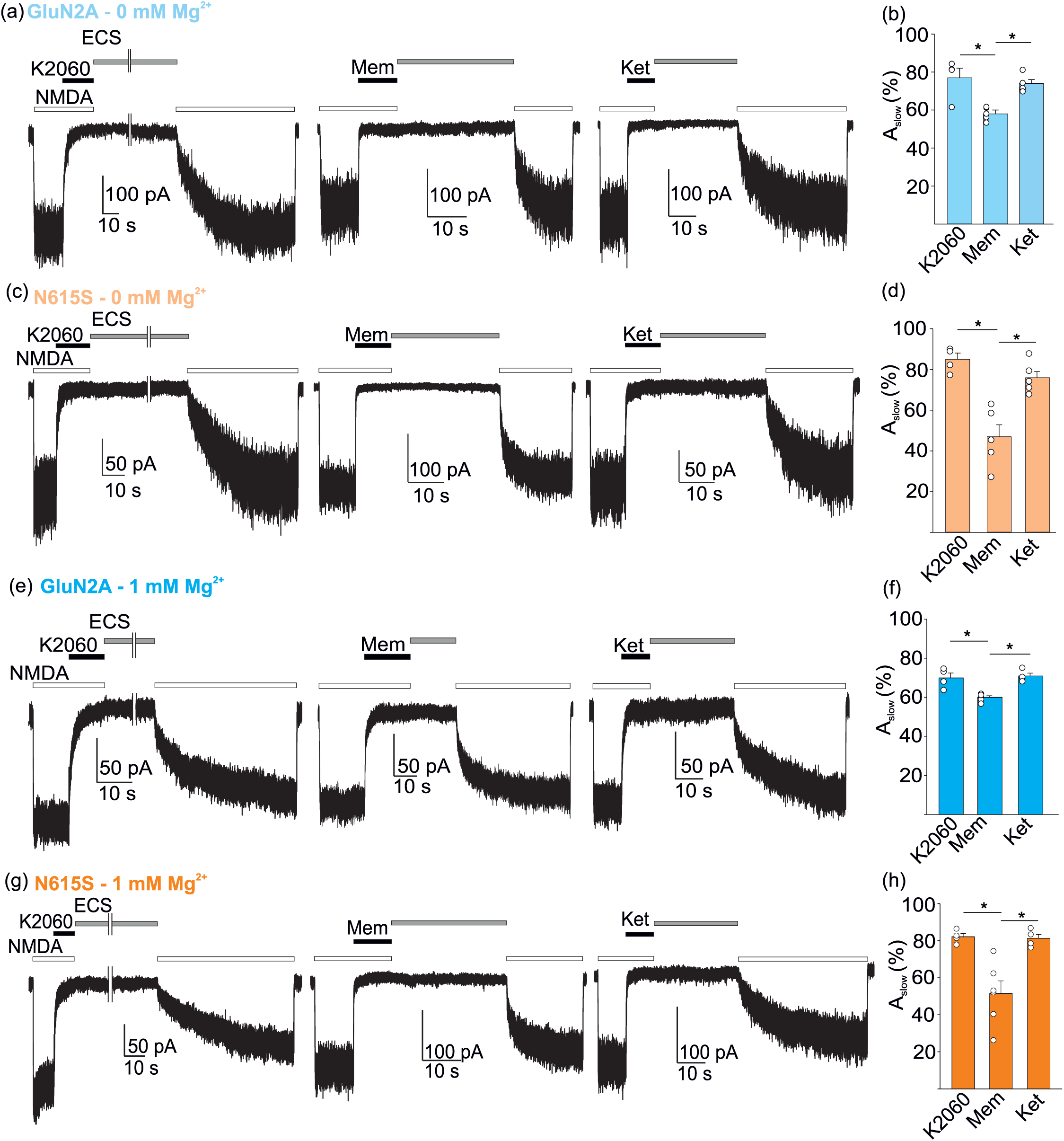
K2060 and ketamine showed greater trapping than memantine independently of the presence of the N615S variant and extracellular Mg^2+^. (a,c,e,g) Representative whole-cell currents recorded from HEK293T cells expressing WT or N615S receptors using a double-pulse protocol to assess recovery from blocker inhibition. GluN1/GluN2A receptors were activated with 30 µM NMDA and 10 µM glycine, followed by application of 100 µM K2060, memantine (Mem) or ketamine (Ket) during their activation. After compound application, cells were washed with extracellular solution (ECS) for a duration corresponding to 10-fold the agonist-present τ_off_ value, and receptors were then reactivated with NMDA and glycine to assess recovery from inhibition. Recordings were performed in the absence or presence of 1 mM Mg^2+^, as indicated. White bars indicate agonist application, black bars indicate compound application, and grey bars indicate ECS washout. Recovery traces were fitted with double-exponential functions; fitted amplitudes and weighted time constants were calculated as described in Methods, with τ_w_ calculated using *Equation 2*. (b,d,f,h) Summary of A_slow_, expressed as a percentage of the total recovery amplitude, obtained from double-exponential fits of recovery from inhibition. A_slow_ was used as an estimate of the fraction of trapped blocker. Data are presented as mean ± SEM, with individual recorded cells shown as open circles. * indicates a statistically significant difference (one-way ANOVA followed by Holm–Sidak all-pairwise *post hoc* test). Exact *n* values and fitted parameters are provided in Supplementary Table S4. Full statistical values are provided in Supplementary Table S7.

### 3.4 Inhibition of endogenous synaptic NMDARs by blockers in acute hippocampal slices exhibited similar patterns to our findings with recombinant NMDARs

After defining potency, Mg^2+^ dependence, kinetics and trapping in recombinant GluN1/GluN2 receptors, we next asked whether the resulting pharmacological profile of the blockers is preserved at synaptic NMDARs under Mg^2+^-containing experimental conditions that closely approximate the *in vivo* extracellular environment. We employed acute hippocampal slices from WT and N615S mice under HEK293T cell-matched recording conditions (internal 10 mM BAPTA; external 1 mM Ca^2+^ and 1 mM Mg^2+^).

To directly compare blocker action at synapses, we evoked pharmacologically isolated NMDAR-eEPSCs in CA1 pyramidal neurons with five pulses at 50 Hz and quantified sustained inhibition over the final 5 min of application of K2060 (Figure 4a), memantine (Figure 4c), and ketamine (Figure 4e). In WT slices, K2060 produced the strongest inhibition, memantine the weakest, with ketamine in between. In N615S slices, inhibition increased for all blockers, with strong inhibition observed for both K2060 and ketamine and lower inhibition by memantine (Figure 4a–g). These results resembled those obtained in HEK293T cells (Figure 4h), although inhibition in WT slices was generally stronger, especially for K2060 and ketamine. These findings indicate that the N615S variant increases sustained inhibition by the blockers at native synaptic NMDARs under Mg^2+^-containing conditions, broadly consistent with the enhanced blocker sensitivity observed with recombinant GluN1/GluN2A-N615S receptors.

**FIGURE 4.**
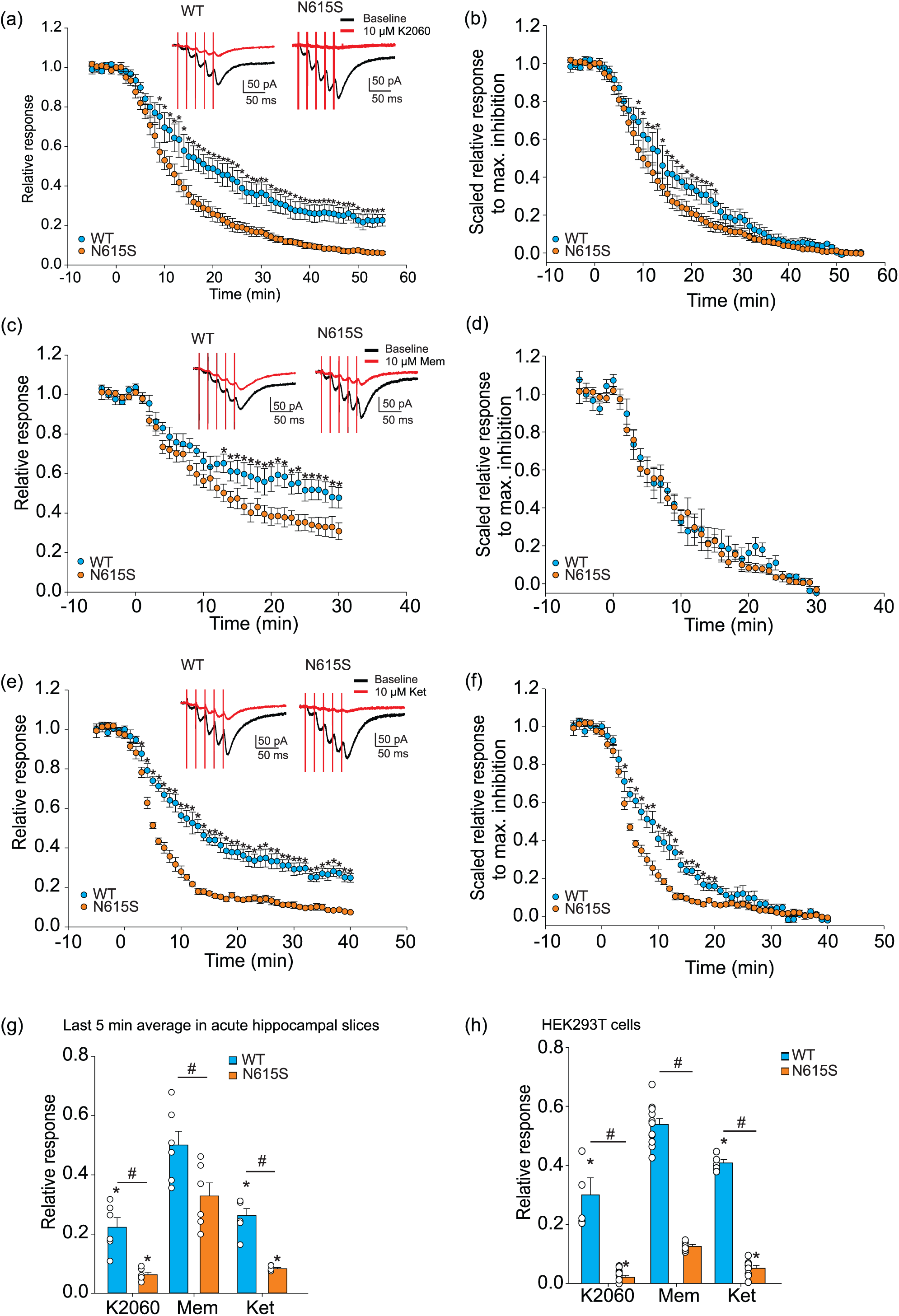
K2060, memantine and ketamine showed enhanced sustained inhibition of native NMDAR-eEPSCs in N615S acute hippocampal slices. (a,c,e) Time course of inhibition of pharmacologically isolated NMDAR-eEPSCs recorded from CA1 pyramidal neurons in acute hippocampal slices from WT and N615S mice. NMDAR-eEPSCs were evoked by trains of five stimuli at 50 Hz in the continuous presence of NBQX and picrotoxin. K2060 (10 µM), memantine (Mem; 10 µM) or ketamine (Ket; 10 µM) was applied after a stable baseline period. Responses were normalised to the mean baseline amplitude before compound application. Insets show representative NMDAR-eEPSCs recorded before blocker application and during the final phase of blocker application. (b,d,f) Same experiments as in (a,c,e), with responses scaled to the maximal inhibition reached during the final 5 min of blocker application to compare inhibition onset kinetics. (g) Mean relative response during the final 5 min of blocker application in acute hippocampal slices (*n*=6 per group). (h) Corresponding comparison of relative responses obtained from HEK293T cells expressing recombinant WT or N615S receptors under extracellular 1 mM Mg^2+^ conditions. Data are presented as mean ± SEM, with individual recorded cells shown as open circles in (g,h). Time-course data in (a–f) were analysed using two-way repeated-measures ANOVA followed by Holm–Sidak *post hoc* test; * indicates a statistically significant difference between WT and N615S at the indicated time point. Summary data in (g,h) were analysed using two-way ANOVA followed by Holm–Sidak *post hoc* test. In (g,h), * indicates a statistically significant difference from memantine within the same genotype or receptor condition; # indicates a statistically significant difference between WT and N615S for the same blocker. Exact *n* values and full statistical values are provided in Supplementary Tables S8 and S9.

To compare the apparent onset of inhibition in slices, we fitted the wash-in phase with a single exponential function to extract τ_on_, and scaled the responses to the mean current amplitude during the final 5 min to visualise the onset kinetics. In WT slices, τ_on_ was ∼19.5 min for K2060, ∼9.8 min for memantine, and ∼11.8 min for ketamine. In N615S slices, the apparent onset was faster for K2060 (∼13.1 min) and ketamine (∼6.2 min), whereas memantine remained similar (∼11.1 min) compared with WT slices. These data aligned with our findings for recombinant NMDARs with K2060 (Table S1), memantine (Table S2), and ketamine (Table S3), while reflecting the more complex nature of hippocampal slice preparation.

### 3.5 N615S mice showed increased behavioural sensitivity to K2060

After confirming that K2060 exhibits enhanced inhibition of N615S receptors expressed in HEK293T cells and in acute slices from N615 mice, we next asked whether this increased pharmacological sensitivity is also reflected *in vivo*. We first used the OFT to assess locomotor activity in WT and N615S mice. Consistent with previous findings (Bertocchi et al., 2021), N615S mice exhibited a hyperlocomotor phenotype (Figure 5a–c). We then administered multiple doses of K2060 (*i.p.*). In WT mice, K2060 did not alter total locomotor activity at doses up to 50 mg/kg (Figure 5d). However, distance travelled did not fully capture dose-related behavioural changes. Mild changes were observed at 30 mg/kg, whereas 50 mg/kg produced slow wall-following, circling-like trajectories, and a sedated appearance. In contrast to WT mice, N615S mice showed greater behavioural sensitivity to K2060 at higher doses. At 5 mg/kg, K2060 had no detectable effect on locomotor behaviour. At 15 mg/kg, K2060 produced mild but noticeable behavioural abnormalities, including corner-directed sniffing, wall-following, mild ataxia and stereotyped movements (Figure 5e,f). These effects were not accompanied by a reduction in total distance travelled, suggesting that locomotor output alone may underestimate behavioural impairment. At 30 mg/kg, K2060 induced a pronounced phenotype characterised by stereotypy, ataxic locomotion, prolonged immobility, and sedation and behavioural arrest-like episodes, resulting in a clear reduction in locomotor activity (Figure 5e,f). These findings showed that N615S mice are more sensitive to K2060 than WT mice. Importantly, 5 mg/kg K2060 did not produce detectable behavioural impairment in the OFT, supporting the use of this dose for subsequent *in vivo* testing.

**FIGURE 5.**
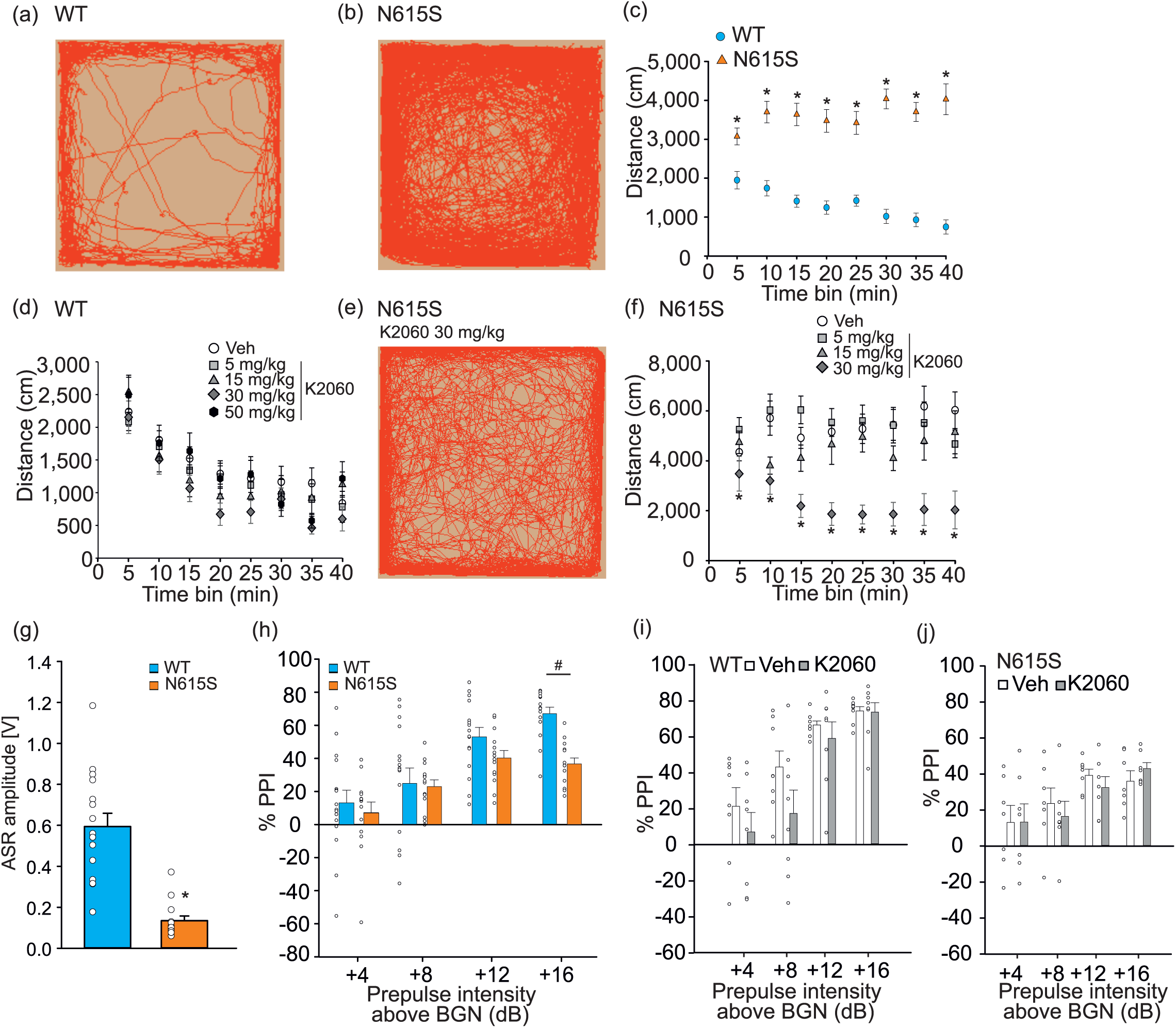
N615S mice showed hyperlocomotion and increased behavioural sensitivity to K2060, whereas 5 mg/kg K2060 did not further disrupt PPI. (a,b) Representative locomotor trajectories recorded during the open OFT in WT (a) and N615S (b) mice under baseline conditions.(c) Distance travelled in 5-min time bins during a 40-min OFT session in WT and N615S mice. N615S mice showed increased locomotor activity compared with WT mice. (d) Distance travelled in 5-min time bins after *i.p.* administration of vehicle (Veh) or K2060 at 5, 15, 30 or 50 mg/kg in WT mice. (e) Representative locomotor trajectory from an N615S mouse treated with K2060 at 30 mg/kg. (f) Distance travelled in 5-min time bins after *i.p.* administration of vehicle or K2060 at 5, 15 or 30 mg/kg in GluN2A-N615S mice. K2060 at 30 mg/kg reduced locomotor activity in N615S mice compared with vehicle. (g) Acoustic startle response (ASR) amplitude in WT and N615S mice. N615S mice showed reduced ASR amplitude compared with WT mice. (h) Baseline prepulse inhibition (PPI) measured at prepulse intensities of +4, +8, +12 and +16 dB above background noise (BGN). N615S mice showed reduced PPI compared with WT mice at higher prepulse intensities. (i,j) PPI measured 30 min after *i.p.* administration of vehicle or K2060 at 5 mg/kg in WT (i) and N615S (j) mice. K2060 did not further disrupt PPI in either genotype. PPI was calculated using *Equation 4*. Data are presented as mean ± SEM, with individual animals shown as open circles where applicable. * indicates a statistically significant difference versus WT at the same time bin or prepulse intensity. # indicates a statistically significant difference versus vehicle or 0 mg/kg K2060 at the same time bin. Statistical analysis was performed using Mann–Whitney rank-sum test for ASR amplitude and two-way repeated-measures ANOVA followed by Holm–Sidak *post hoc* test where appropriate. Exact *n* values and full statistical details are provided in Supplementary Table S10-11.

### 3.6 K2060 did not further disrupt PPI in WT or N615S mice

Because the OFT primarily measures locomotor activity and may not reveal changes in sensorimotor gating, we next performed a focused ASR/PPI experiment with K2060 at 5 mg/kg (*i.p.*). N615S mice showed markedly weaker startle reactivity than WT littermates, with a ∼5-fold lower mean ASR amplitude (Figure 5g). PPI was also lower in N615S than WT mice at higher prepulse intensities, indicating impaired sensorimotor gating (Figure 5h). We then tested whether administration of vehicle or K2060 at 5 mg/kg (*i.p.*) alters PPI. Our data showed that PPI did not differ between treatment groups in either WT or N615S mice (Figure 5i,j). Thus, although N615S mice displayed baseline deficits in ASR and PPI, acute K2060 treatment at 5 mg/kg (*i.p.*) did not further disrupt PPI.

### 3.7 K2060 protected N615S mice against AGS at a lower dose than memantine or ketamine

Having established that 5 mg/kg (*i.p.*) K2060 did not produce detectable behavioural impairment in the OFT or further disrupt PPI, we next tested whether this dose protects N615S mice against AGS. Building on prior work showing memantine (5 mg/kg, *i.p.*) prevents AGS in N615S mice (Bertocchi et al., 2021), we compared K2060 with memantine and ketamine at the same dose and route of administration. Male N615S mice received K2060, memantine, ketamine, or vehicle and, 30 min later, were exposed to acoustic stimulation until AGS occurred or the protocol ended. Vehicle-treated mice showed mostly AGS with and without RA (Figure 6a). At 5 mg/kg (*i.p.*), all three blockers were effective, providing protection in most animals. To discriminate among blockers, we reduced the dose of all three blockers to 1 mg/kg (*i.p.*; Figure 6b). Then, we observed that only K2060 protected N615S mice compared with vehicle, whereas memantine and ketamine did not. These findings showed that K2060 exhibits favourable *in vivo* potency to protect against AGS in N615S mice.

**FIGURE 6.**
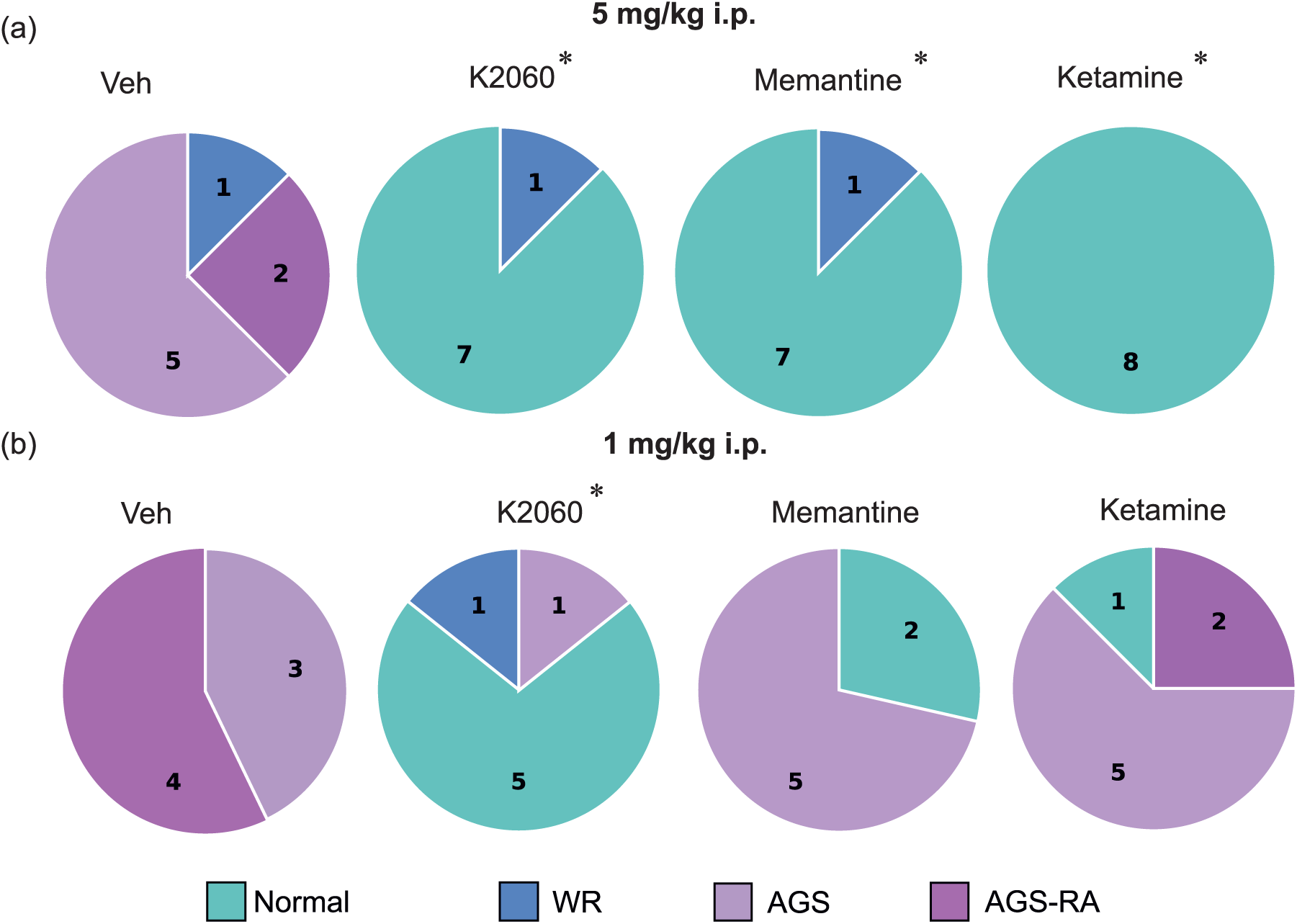
K2060 protected N615S mice against audiogenic seizures at a lower dose than memantine or ketamine. Male N615S mice were treated *i.p.* with vehicle (Veh), K2060, memantine or ketamine at 5 (a) or 1 (b) mg/kg 30 min before acoustic stimulation. Pie charts show the proportion of animals classified as normal response, wild running (WR), audiogenic seizure (AGS) or AGS followed by respiratory arrest (AGS-RA). Numbers inside pie charts indicate the number of animals in each outcome category. At 5 mg/kg, all three blockers reduced AGS severity compared with vehicle. At 1 mg/kg, only K2060 significantly protected N615S mice compared with the corresponding vehicle group. Statistical analysis was performed using Fisher’s exact test. * indicates a statistically significant difference versus the corresponding dose-matched vehicle group. Numbers inside pie charts indicate the number of animals in each outcome category. Full statistical details are provided in Supplementary Table S11.

### 3.8 K2060 showed a similar pharmacological profile at N615S and GluN1/GluN2A-N615K (N615K) receptors

Finally, we asked whether the pharmacological profile observed at N615S receptors also extends to recombinant GluN1/GluN2A receptors carrying the pathogenic N615K variant (N615K receptors), identified in patients with severe neurodevelopmental phenotypes (Endele et al., 2010; Allen et al., 2016; Li et al., 2019). First, concentration–response analysis with Mg^2+^ confirmed reduced Mg^2+^ block at N615K receptors, consistent with prior work (Endele et al., 2010) (Figure 7a,b; Table 1). We then performed concentration–response analysis with all three blockers at N615K receptors. K2060 and ketamine showed similar apparent potencies at N615K and N615S receptors, whereas memantine was ∼3.5-fold less potent at N615K receptors. The presence of Mg^2+^ did not substantially alter IC_50_ values of all three blockers at N615K receptors, similarly to what we observed for N615S receptors (Figure 7b; Table 1).

**FIGURE 7.**
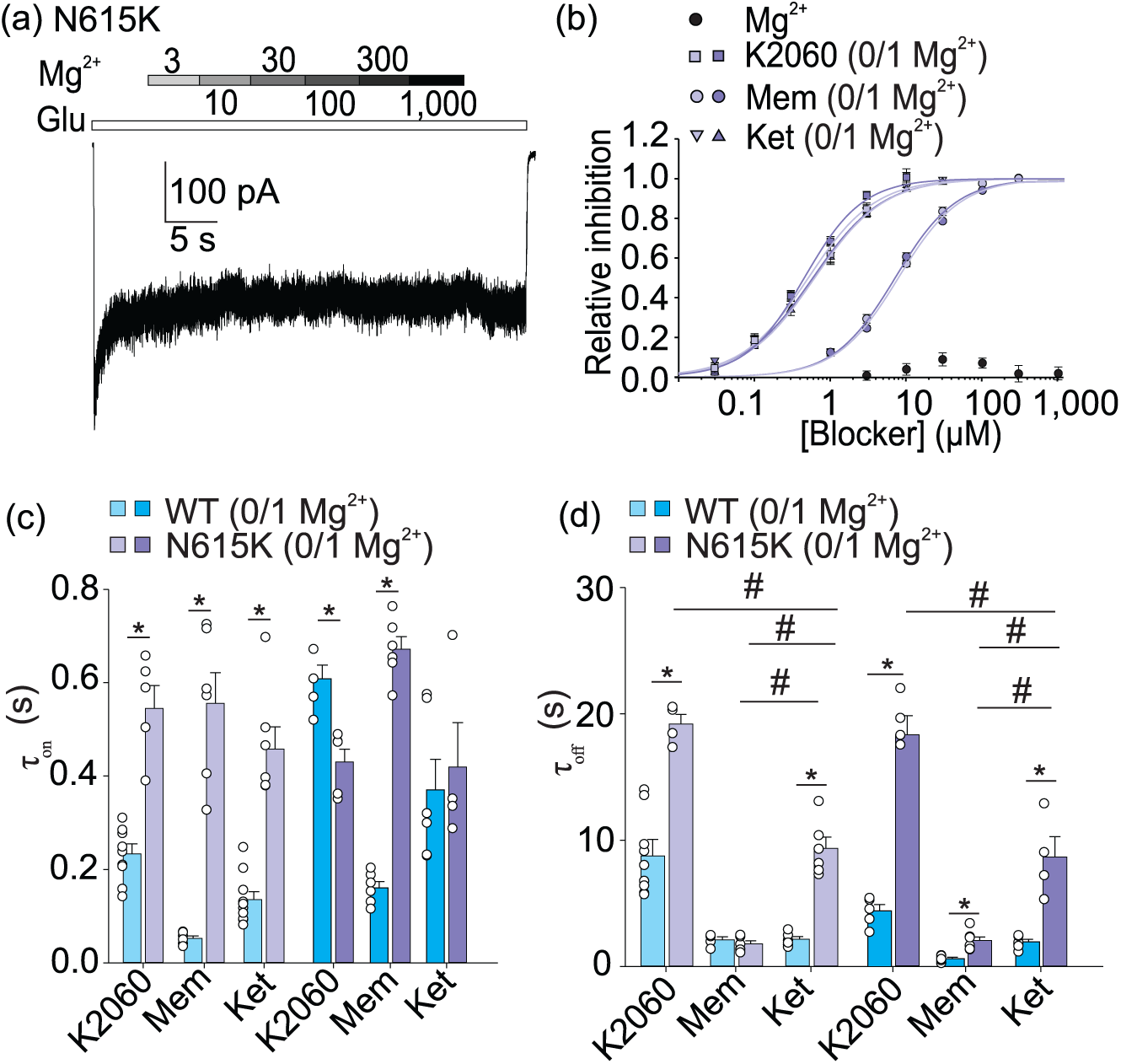
Inhibition of N615K receptors by memantine, ketamine and K2060. (a) Representative whole-cell current responses recorded from a HEK293T cell expressing N615K receptors during application of increasing concentrations of extracellular Mg^2+^. Current responses were evoked by 1 mM glutamate (Glu; white bar) in the continuous presence of 50 µM glycine. (b) Concentration-response relationships for inhibition of N615K receptors by Mg^2+^ and by memantine (Mem), ketamine (Ket) or K2060. Blocker concentration-response relationships were obtained in the absence and presence of 1 mM extracellular Mg^2+^. Data are presented as mean ± SEM. Concentration-response data were fitted with *Equation 1*. Fitted IC_50_ values and Hill coefficients are provided in Table 1; full statistical details are provided in Supplementary Table S5. (c,d) Summary of onset (τ_on_; c) and recovery (τ_off_; d) time constants for inhibition by 30 µM memantine, ketamine or K2060 at WT and N615K receptors in the absence or presence of 1 mM Mg^2+^. Kinetic time constants were obtained as described in Methods, with weighted recovery time constants calculated using *Equation 2*. Data are presented as mean ± SEM, with individual recorded cells shown as open circles. * indicates a statistically significant difference between WT and N615K receptors for the same blocker and Mg^2+^ condition. # indicates a statistically significant difference between the indicated 0 and 1 mM Mg^2+^ conditions. Statistical comparisons were performed as described in Methods. Exact *n* values and kinetic parameters for c,d are provided in Supplementary Tables S1–S3.

Using saturating concentrations of the blockers (30 μM), we next analysed τ_on_ and τ_off_ in the presence of 0 and 1 mM Mg^2+^ (Figure 7c,d; Tables S1–3). At N615K receptors, τ_on_ was similar across blockers and Mg^2+^ conditions. Compared with N615S receptors, τ_on_ was largely unchanged for K2060, whereas it was ∼10-fold slower for memantine and ∼5-fold slower for ketamine. Concerning τ_off_, K2060 and memantine showed comparable values at N615K and N615S receptors, whereas ketamine recovery was markedly prolonged at N615K receptors. These data showed that K2060 maintains similar apparent potency and kinetic profiles at both N615S and N615K receptors.

## 4 DISCUSSION

We focused on K2060, a dizocilpine-derived compound recently characterised as a potent, reversible NMDAR blocker (Misiachna et al., 2024) and compared its inhibitory action with the clinically relevant blockers memantine and ketamine. Across preparations, extracellular Mg^2+^ strongly modulated blocker potency and kinetics at the WT NMDARs. In HEK293T cells, the N615S variant strongly reduced Mg^2+^ block, consistent with previous work (Wollmuth et al., 1998; Bertocchi et al., 2021). In the absence of extracellular Mg^2+^, IC_50_ values for memantine, ketamine, and K2060 were similar between WT and N615S receptors, suggesting that their binding sites within the ion channel pore are preserved. In 1 mM Mg^2+^, only WT receptors showed the expected rightward potency shift of all studied blockers. This indicates that impaired Mg^2+^ block removes a physiological constraint on blocker inhibition of NMDARs and supports the hypothesis that Mg^2+^ competes with blockers within the ion channel pore.

The kinetic data further support this interpretation. Mg^2+^ slowed τ_on_ at WT but not at N615S receptors (Sobolevsky et al., 1998; Kashiwagi et al., 2002; Kotermanski and Johnson, 2009; Glasgow et al., 2018; Kolcheva et al., 2021, 2023). The N615S-specific slowing of K2060 kinetics suggests that the substitution at the 615 residue, within the M2-loop selectivity filter, perturbs local ion channel pore architecture in a way that selectively affects this chemotype, consistent with heterogeneous blocker–ion channel pore interactions across different NMDAR ligands (Chou et al., 2022; Zhou and Tajima, 2023). Mg^2+^ also accelerated τ_off_ for memantine and K2060 at WT but not N615S receptors, in line with previous data (Glasgow et al., 2018). Thus, Mg^2+^ shapes block at WT receptors, whereas the N615S variant abolishes this effect.

We observed partial trapping for all three blockers, ranked memantine < ketamine ≤ K2060, in agreement with prior reports for memantine (Blanpied et al., 1997; Kotermanski et al., 2009), near-complete trapping for ketamine (Mealing et al., 1999; Bettini et al., 2022; Ma et al., 2023), and strong trapping for K2060 (Misiachna et al., 2024). The degree of trapping was similar at WT and N615S receptors and remained unchanged in the presence of Mg^2+^, indicating that it is governed by intrinsic blocker–ion-channel pore determinants. This is supported by the fact that ketamine is less lipophilic than memantine yet more strongly trapped within the NMDARs (Mealing et al., 2001; Bolshakov et al., 2003).

We next examined whether these pharmacological differences were preserved in a neuronal preparation. In our hippocampal slice experiments, we used 5 pulses at 50 Hz every 20 s to achieve stable blocker inhibition because use-dependent block scales with stimulation frequency (Wild et al., 2013) and reaching a single-pulse steady-state level can take hours (Wallach et al., 2016). In WT slices, steady-state inhibition levels resembled data from the HEK293T cells, but their degree was generally stronger, especially for ketamine and K2060, which we attribute to greater blocker trapping during brief, burst-like synaptic openings (consistent with trapping studied in the HEK293T cells; Figure 2; Table S4). In N615S slices, steady-state degree of blocker inhibition was lower than observed in the HEK293T cells, likely reflecting NMDAR heterogeneity, including GluN1/GluN2B and GluN1/GluN2A/GluN2B receptors (Zhang et al., 2025). Concerning blocker kinetics, HEK293T cells-to-slice trends were largely preserved as memantine showed similar τ_on_ in slices despite faster onset at N615S receptors in HEK293T cells, likely due to prolonged τ_off_; ketamine displayed a clear acceleration of τ_on_ in N615S slices, consistent with faster τ_on_ and only minor τ_off_ changes at recombinant receptors in HEK293T cells. For K2060, the markedly prolonged τ_off_ at N615S receptors expressed in HEK293T cells likely contributes to the faster apparent onset observed in N615S slices. We cannot rule out the possibility that additional factors, such as membrane-to-channel access (Neureiter et al., 2025), may also contribute.

The *in vivo* experiments were designed first to assess the tolerability of K2060 over a broad dose range. Acute ketamine administration has been reported to increase locomotor activity in rodents at doses of ∼25–75 mg/kg (*i.p.*). Ketamine-induced behavioural activation may also be accompanied by stereotyped behaviour and ataxia, while higher doses can produce an initial suppression of locomotion followed by delayed hyperactivity (Razoux et al., 2007; Caixeta et al., 2013; Chen et al., 2023). Moreover, behavioural responses to ketamine can differ among C57BL/6 mouse substrains, although evidence from subchronic administration cannot be directly extrapolated to acute locomotor responses (Harda et al., 2022). Acute memantine administration has likewise been reported to induce hyperlocomotion in mice at doses of 10–20 mg/kg (*i.p.*), with the higher dose also producing persistent stereotyped behaviour (Kitanaka et al., 2018; Ueno et al., 2020). Thus, the observed locomotor response likely depends not only on animal species and dose, but also on the timing and duration of behavioural assessment, because motor impairment or stereotypies may initially mask locomotor activation and the hyperlocomotor component may emerge later (Liljequist, 1991; Wu et al., 2005; Qi et al., 2008).

In our experiments, K2060 did not reduce total distance travelled at doses up to 50 mg/kg (*i.p.*) in WT mice, suggesting its safety compared to ketamine and memantine. In contrast, in N615S mice, K2060 reduced total distance travelled at doses from 30 mg/kg (*i.p.*); in addition, K2060 exhibited more pronounced behavioural side effects. These findings suggest increased *in vivo* sensitivity to K2060 under impaired Mg^2+^ block of the NMDARs. Furthermore, we observed that K2060 did not disrupt PPI in either WT or N615S mice. Previous studies have shown that memantine has a dose-dependent effect on PPI; at 5 mg/kg (*i.p.*), it can produce a slight disruption (Nakaya et al., 2011), and ketamine similarly induces PPI deficits at 5 mg/kg, but not at 1 mg/kg (both *i.p.*), in rats (Kamiyama et al., 2011). However, such comparisons should also be interpreted with caution because PPI effects may be influenced by the experimental protocol, animal species, and substrain. For example, C57BL/6 mouse substrains (B6N vs B6J) differ in PPI, and ddY mice can show markedly different responses from C57BL/6 mice (Simon et al., 2013). In addition, mouse–rat comparisons are not straightforward, as rats often require ∼2-fold higher doses to elicit comparable effects (Nair and Jacob, 2016).

Previous studies have shown that N615S mice are highly susceptible to AGS and that this phenotype can be rescued by NMDAR-targeted pharmacological interventions, including memantine and radiprodil (Bertocchi et al., 2021, 2024). Our data extended these findings by showing that K2060, memantine and ketamine all protected these mice at 5 mg/kg (*i.p.*), whereas only K2060 remained protective at 1 mg/kg (*i.p.*). As memantine has favourable brain exposure and persistence (Beconi et al., 2011), and ketamine appears broadly comparable to K2060 in pharmacokinetic terms (Toki et al., 2018), this suggests that K2060’s efficacy reflects ion-channel-blocking properties rather than a simple pharmacokinetic advantage.

Finally, we examined whether the pharmacological effects observed for N615S receptors extended to the disease-associated N615K receptors. Loss of Mg^2+^ block can increase NMDAR signalling at negative membrane potentials and contribute to *GRIN*-associated neurodevelopmental and epileptic phenotypes (Wollmuth et al., 1998; Endele et al., 2010; Bertocchi et al., 2021; Santos-Gómez et al., 2021). We employed N615S mice as an *in vivo* model of impaired Mg^2+^ block, although the N615S variant has not been reported to be pathogenic. In contrast, the N615K variant has been linked to severe neurodevelopmental phenotypes, including developmental delay, intellectual disability, hypotonia, language impairment, behavioural anomalies, and multiple epilepsy types (Endele et al., 2010; Allen et al., 2016; Li et al., 2019). Both variants reduced Mg^2+^ sensitivity, but they differed in how they shaped blocker action. N615K receptors showed substantially reduced memantine potency, whereas K2060 and ketamine potencies were largely preserved, consistent with *Xenopus* oocyte data showing impaired memantine inhibition but only slight alterations in ketamine inhibition at N615K receptors (Marwick et al., 2015, 2019). N615K receptors also strongly slowed the τ_on_ relative to N615S receptors, especially for memantine and ketamine, and markedly prolonged ketamine τ_off_. In contrast, K2060 maintained similar potency and kinetic profile across N615S and N615K receptors. These differences likely reflect residue-specific blocker–ion channel pore interactions and/or alterations of the single-channel conductance (Marwick et al., 2019).

## 5 CONCLUSION

Impaired Mg^2+^ block of GluN1/GluN2A receptors caused by substitutions of the GluN2A-N615 selectivity-filter residue reshapes the potency and kinetics of blockers, including memantine, ketamine and K2060. K2060 retained strong inhibitory activity at the mutated NMDARs and protected N615S mice against AGS at a lower dose than memantine or ketamine. Therefore, K2060 is a promising compound for further preclinical development in NMDAR hyperfunction states, specifically those associated with pathogenic missense variants in the ion channel pore region that impair Mg^2+^ block. Further pharmacokinetic, safety, and broader disease-model studies remain necessary.

## Acknowledgments

This work was supported by the Czech Science Foundation (24-10026S; supporting studies of K2060), by the Ministry of Education, Youth and Sports of the Czech Republic (ExRegMed CZ.02.01.01/00/22_008/0004562; supporting experiments with memantine), and by the Ministry of Health of the Czech Republic in cooperation with the Czech Health Research Council (No. NW24J-04-00084; supporting studies involving ketamine). We thank Rolf Sprengel for providing the mutant mouse line (Allele: Grin2a<tm2Rsp>), INFRAFRONTIER/EMMA (www.infrafrontier.eu, NFRAFRONTIER Consortium, 2015), and the EMMA node at Karolinska Institute from which the mouse line was distributed (RRID:IMSR_EM:09319).

## AUTHOR CONTRIBUTIONS

Conceptualisation: Marharyta Kolcheva, Marek Ladislav and Martin Horak. Methodology: Marharyta Kolcheva, Jakub Netolicky, Jan Korabecny, Ondrej Soukup, Marek Ladislav and Martin Horak. Investigation: Marharyta Kolcheva, Jakub Netolicky, Anna Misiachna, Natalia Rybalkova, Petra Zahumenska, Jovana Doderovic, Alena Pakhomova, Claudia Valentina De Sanctis, Gregory Klein Schneider, Katarina Hemelikova, Kristyna Rehakova, Rostislav Turecek. Formal analysis: Marharyta Kolcheva, Jakub Netolicky and Anna Misiachna. Data curation: Marharyta Kolcheva, Marek Ladislav and Martin Horak. Writing original draft: Marharyta Kolcheva, Marek Ladislav, Ondrej Soukup, Martin Horak. Visualisation: Marharyta Kolcheva, Jakub Netolicky and Marek Ladislav. Supervision: Marek Ladislav and Martin Horak. Project administration: Marek Ladislav and Martin Horak.

## DATA AVAILABILITY STATEMENT

The data supporting the findings are available from Zenodo at DOI: 10.5281/zenodo.20797987.

## DECLARATION OF TRANSPARENCY AND SCIENTIFIC RIGOUR

This Declaration acknowledges that this paper adheres to the principles for transparent reporting and scientific rigour of preclinical research as stated in the *BJP* guidelines for Design and Analysis and Animal Experimentation, and as recommended by funding agencies, publishers, and other organisations engaged with supporting research.

## Abbreviations

aCSF: artificial cerebrospinal fluid
AGS: audiogenic seizures
AGS-RA: AGS followed by respiratory arrest
ASR: acoustic startle response
DMSO: dimethyl sulfoxide
ECS: extracellular solution
IC_50_: half-maximal inhibitory concentration
ICS: intracellular solution
NMDAR: N-methyl-D-aspartate receptor
NMDAR-eEPSCs: evoked NMDAR-mediated excitatory postsynaptic currents
OFT: open field test
PPI: prepulse inhibition
RA: respiratory arrest
WR: wild running
WT: wild-type.

## REFERENCES

Alexander, S. P. H., Davenport, A. P., Kelly, E., Gibb, A. J., Mathie, A. A., Peach, C. J., et al. (2025). The Concise Guide to PHARMACOLOGY 2025/26: G protein-coupled receptors. British Journal of Pharmacology, 182. 10.1111/bph.70230

Allen, N. M., Conroy, J., Shahwan, A., Lynch, B., Correa, R. G., Pena, S. D. J., et al. (2016). Unexplained early onset epileptic encephalopathy: Exome screening and phenotype expansion. Epilepsia, 57. 10.1111/epi.13250

Beconi, M. G., Howland, D., Park, L., Lyons, K., Giuliano, J., Dominguez, C., et al. (2011). Pharmacokinetics of memantine in rats and mice. PLoS Currents, 3, RRN1291. 10.1371/currents.rrn1291

Benke, T. A., Park, K., Krey, I., Camp, C. R., Song, R., Ramsey, A. J., et al. (2021). Clinical and therapeutic significance of genetic variation in the GRIN gene family encoding NMDARs. Neuropharmacology, 199, 108805. 10.1016/j.neuropharm.2021.108805

Bertocchi, I., Cifarelli, L., Oberto, A., Eva, C. E., Sprengel, R., Mirza, N. R., et al. (2024). Radiprodil, a selective GluN2B negative allosteric modulator, rescues audiogenic seizures in mice carrying the GluN2A(N615S) mutation. British Journal of Pharmacology, 181, 1886–1894. 10.1111/bph.16361

Bertocchi, I., Eltokhi, A., Rozov, A., Chi, V. N., Jensen, V., Bus, T., et al. (2021). Voltage-independent GluN2A-type NMDA receptor Ca2+ signaling promotes audiogenic seizures, attentional and cognitive deficits in mice. Communications Biology, 4, 59. 10.1038/s42003-020-01538-4

Bettini, E., Stahl, S. M., De Martin, S., Mattarei, A., Sgrignani, J., Carignani, C., et al. (2022). Pharmacological Comparative Characterization of REL-1017 (Esmethadone-HCl) and Other NMDAR Channel Blockers in Human Heterodimeric N-Methyl-D-Aspartate Receptors. Pharmaceuticals, 15, 997. 10.3390/ph15080997

Blanpied, T. A., Boeckman, F. A., Aizenman, E., & Johnson, J. W. (1997). Trapping Channel Block of NMDA-Activated Responses By Amantadine and Memantine. Journal of Neurophysiology, 77, 309–323. 10.1152/jn.1997.77.1.309

Bolshakov, K. V., Gmiro, V. E., Tikhonov, D. B., & Magazanik, L. G. (2003). Determinants of trapping block of N-methyl-d-aspartate receptor channels. Journal of Neurochemistry, 87, 56–65. 10.1046/j.1471-4159.2003.01956.x

Caixeta, F. V., Cornélio, A. M., Scheffer-Teixeira, R., Ribeiro, S., & Tort, A. B. L. (2013). Ketamine alters oscillatory coupling in the hippocampus. Scientific Reports, 3, 2348. 10.1038/srep02348

Chen, W.-C., Wang, T.-S., Chang, F.-Y., Chen, P.-A., & Chen, Y.-C. (2023). Age, Dose, and Locomotion: Decoding Vulnerability to Ketamine in C57BL/6J and BALB/c Mice. Biomedicines, 11, 1821. 10.3390/biomedicines11071821

Chou, T.-H., Epstein, M., Michalski, K., Fine, E., Biggin, P. C., & Furukawa, H. (2022). Structural insights into binding of therapeutic channel blockers in NMDA receptors. Nature Structural & Molecular Biology, 29, 507–518. 10.1038/s41594-022-00772-0

Curtis, M. J., Alexander, S. P. H., Cirino, G., George, C. H., Kendall, D. A., Insel, P. A., et al. (2022). Planning experiments: Updated guidance on experimental design and analysis and their reporting III. British Journal of Pharmacology, 179, 3907–3913. 10.1111/bph.15868

Endele, S., Rosenberger, G., Geider, K., Popp, B., Tamer, C., Stefanova, I., et al. (2010). Mutations in GRIN2A and GRIN2B encoding regulatory subunits of NMDA receptors cause variable neurodevelopmental phenotypes. Nature Genetics, 42, 1021–1026. 10.1038/ng.677

Fedele, L., Newcombe, J., Topf, M., Gibb, A., Harvey, R. J., & Smart, T. G. (2018). Disease-associated missense mutations in GluN2B subunit alter NMDA receptor ligand binding and ion channel properties. Nature Communications, 9, 957. 10.1038/s41467-018-02927-4

Glasgow, N. G., Wilcox, M. R., & Johnson, J. W. (2018). Effects of Mg2+ on recovery of NMDA receptors from inhibition by memantine and ketamine reveal properties of a second site. Neuropharmacology, 137, 344–358. 10.1016/j.neuropharm.2018.05.017

Hansen, K. B., Wollmuth, L. P., Bowie, D., Furukawa, H., Menniti, F. S., Sobolevsky, A. I., et al. (2021). Structure, Function, and Pharmacology of Glutamate Receptor Ion Channels. Pharmacological Reviews, 73, 1469–1658. 10.1124/pharmrev.120.000131

Harda, Z., Misiołek, K., Klimczak, M., Chrószcz, M., & Rodriguez Parkitna, J. (2022). C57BL/6N mice show a sub-strain specific resistance to the psychotomimetic effects of ketamine. Frontiers in Behavioral Neuroscience, 16, 1057319. 10.3389/fnbeh.2022.1057319

Harding, S. D., Armstrong, J. F., Faccenda, E., Southan, C., Alexander, S. P. H., Davenport, A. P., et al. (2026). The IUPHAR/BPS Guide to PHARMACOLOGY in 2026. Nucleic Acids Research, 54, D1446–D1456. 10.1093/nar/gkaf1067

Huang, X., Sun, X., Wang, Q., Zhang, J., Wen, H., Chen, W.-J., et al. (2025). Structural insights into the diverse actions of magnesium on NMDA receptors. Neuron, 113, 1006–1018.e4. 10.1016/j.neuron.2025.01.021

INFRAFRONTIER Consortium (2015). INFRAFRONTIER--providing mutant mouse resources as research tools for the international scientific community. Nucleic Acids Research, 43, D1171– D1175. 10.1093/nar/gku1193

Kamiyama, H., Matsumoto, M., Otani, S., Kimura, S.-I., Shimamura, K.-I., Ishikawa, S., et al. (2011). Mechanisms underlying ketamine-induced synaptic depression in rat hippocampus-medial prefrontal cortex pathway. Neuroscience, 177, 159–169. 10.1016/j.neuroscience.2010.12.012

Kashiwagi, K., Masuko, T., Nguyen, C. D., Kuno, T., Tanaka, I., Igarashi, K., et al. (2002). Channel Blockers Acting at N-Methyl-d-aspartate Receptors: Differential Effects of Mutations in the Vestibule and Ion Channel Pore. Molecular Pharmacology, 61, 533–545. 10.1016/s0026-895x(24)12116-5

Kitanaka, N., Kitanaka, J., Hall, F. S., Kubota, Y., Mimura, Y., Ogura, S., et al. (2018). Psychotomimetic-like behavioral effects of memantine in the mouse. Biomedicine & Pharmacotherapy, 100, 116–123. 10.1016/j.biopha.2018.01.160

Kolcheva, M., Kortus, S., Krausova, B. H., Barackova, P., Misiachna, A., Danacikova, S., et al. (2021). Specific pathogenic mutations in the M3 domain of the GluN1 subunit regulate the surface delivery and pharmacological sensitivity of NMDA receptors. Neuropharmacology, 189, 108528. 10.1016/j.neuropharm.2021.108528

Kolcheva, M., Ladislav, M., Netolicky, J., Kortus, S., Rehakova, K., Krausova, B. H., et al. (2023). The pathogenic N650K variant in the GluN1 subunit regulates the trafficking, conductance, and pharmacological properties of NMDA receptors. Neuropharmacology, 222, 109297. 10.1016/j.neuropharm.2022.109297

Korinek, M., Candelas Serra, M., Abdel Rahman, F. E. S., Dobrovolski, M., Kuchtiak, V., Abramova, V., et al. (2024). Disease-Associated Variants in GRIN1, GRIN2A and GRIN2B genes: Insights into NMDA Receptor Structure, Function, and Pathophysiology. Physiological Research, S413–S434. 10.33549/physiolres.935346

Kotermanski, S. E., & Johnson, J. W. (2009). Mg2+Imparts NMDA Receptor Subtype Selectivity to the Alzheimer’s Drug Memantine. The Journal of Neuroscience, 29, 2774–2779. 10.1523/jneurosci.3703-08.2009

Kotermanski, S. E., Wood, J. T., & Johnson, J. W. (2009). Memantine binding to a superficial site on NMDA receptors contributes to partial trapping. The Journal of Physiology, 587, 4589–4604. 10.1113/jphysiol.2009.176297

Li, J., Zhang, J., Tang, W., Mizu, R. K., Kusumoto, H., XiangWei, W., et al. (2019). De novoGRINvariants in NMDA receptor M2 channel pore-forming loop are associated with neurological diseases. Human Mutation, 40, 2393–2413. 10.1002/humu.23895

Liljequist, S. (1991). Genetic differences in the effects of competitive and non-competitive NMDA receptor antagonists on locomotor activity in mice. Psychopharmacology, 104, 17–21. 10.1007/bf02244548

Lilley, E., Stanford, S. C., Kendall, D. E., Alexander, S. P., Cirino, G., Docherty, J. R., et al. (2020). ARRIVE 2.0 and the British Journal of Pharmacology: Updated guidance for 2020. British Journal of Pharmacology, 177, 3611–3616. 10.1111/bph.15178

Ma, S., Chen, M., Jiang, Y., Xiang, X., Wang, S., Wu, Z., et al. (2023). Sustained antidepressant effect of ketamine through NMDAR trapping in the LHb. Nature, 622, 802–809. 10.1038/s41586-023-06624-1

Marwick, K., Skehel, P., Hardingham, G., & Wyllie, D. (2015). Effect of a GRIN2A de novo mutation associated with epilepsy and intellectual disability on NMDA receptor currents and Mg2+ block in cultured primary cortical neurons. The Lancet, 385, S65. 10.1016/s0140-6736(15)60380-4

Marwick, K. F. M., Skehel, P. A., Hardingham, G. E., & Wyllie, D. J. A. (2019). The human NMDA receptor GluN2AN615K variant influences channel blocker potency. Pharmacology Research & Perspectives, 7, e00495. 10.1002/prp2.495

Mealing, G., Lanthorn, T., Murray, C., Small, D., & Morley, P. (1999). Differences in Degree of Trapping of Low-Affinity Uncompetitive N-Methyl-d-aspartic Acid Receptor Antagonists with Similar Kinetics of Block. The Journal of Pharmacology and Experimental Therapeutics, 288, 204–210. 10.1016/s0022-3565(24)37943-1

Mealing, G., Lanthorn, T., Small, D., Murray, R., Mattes, K., Comas, T., et al. (2001). Structural Modifications to anN-Methyl-d-aspartate Receptor Antagonist Result in Large Differences in Trapping Block. The Journal of Pharmacology and Experimental Therapeutics, 297, 906–914. 10.1016/s0022-3565(24)29614-2

Misiachna, A., Konecny, J., Kolcheva, M., Ladislav, M., Prchal, L., Netolicky, J., et al. (2024). Potent and reversible open-channel blocker of NMDA receptor derived from dizocilpine with enhanced membrane-to-channel inhibition. Biomedicine & Pharmacotherapy, 178, 117201. 10.1016/j.biopha.2024.117201

Monyer, H., Sprengel, R., Schoepfer, R., Herb, A., Higuchi, M., Lomeli, H., et al. (1992). Heteromeric NMDA Receptors: Molecular and Functional Distinction of Subtypes. Science, 256, 1217–1221. 10.1126/science.256.5060.1217

Nair, A., & Jacob, S. (2016). A simple practice guide for dose conversion between animals and human. Journal of Basic and Clinical Pharmacy, 7, 27. 10.4103/0976-0105.177703

Neureiter, E. G., Erickson-Oberg, M. Q., Nigam, A., & Johnson, J. W. (2025). Inhibition of NMDA receptors and other ion channel types by membrane-associated drugs. Frontiers in Pharmacology, 16, 1561956. 10.3389/fphar.2025.1561956

Paoletti, P., Bellone, C., & Zhou, Q. (2013). NMDA receptor subunit diversity: impact on receptor properties, synaptic plasticity and disease. Nature Reviews Neuroscience, 14, 383–400. 10.1038/nrn3504

Percie du Sert, N., Hurst, V., Ahluwalia, A., Alam, S., Avey, M. T., Baker, M., et al. (2020). The ARRIVE guidelines 2.0: Updated guidelines for reporting animal research. PLOS Biology, 18, e3000410. 10.1371/journal.pbio.3000410

Platzer, K., Yuan, H., Schütz, H., Winschel, A., Chen, W., Hu, C., et al. (2017). GRIN2B encephalopathy: novel findings on phenotype, variant clustering, functional consequences and treatment aspects. Journal of Medical Genetics, 54, 460–470. 10.1136/jmedgenet-2016-104509

Qi, C., Zou, H., Zhang, R., Zhao, G., Jin, M., & Yu, L. (2008). Age-related differential sensitivity to MK-801-induced locomotion and stereotypy in C57BL/6 mice. European Journal of Pharmacology, 580, 161–168. 10.1016/j.ejphar.2007.07.071

Razoux, F., Garcia, R., & Léna, I. (2007). Ketamine, at a Dose that Disrupts Motor Behavior and Latent Inhibition, Enhances Prefrontal Cortex Synaptic Efficacy and Glutamate Release in the Nucleus Accumbens. Neuropsychopharmacology, 32, 719–727. 10.1038/sj.npp.1301057

Santos-Gómez, A., Miguez-Cabello, F., Juliá-Palacios, N., García-Navas, D., Soto-Insuga, V., García-Peñas, J. J., et al. (2021). Paradigmatic De Novo GRIN1 Variants Recapitulate Pathophysiological Mechanisms Underlying GRIN1-Related Disorder Clinical Spectrum. International Journal of Molecular Sciences, 22, 12656. 10.3390/ijms222312656

Seibenhener, M. L., & Wooten, M. C. (2015). Use of the Open Field Maze to Measure Locomotor and Anxiety-like Behavior in Mice. Journal of Visualized Experiments. 10.3791/52434

Simon, M. M., Greenaway, S., White, J. K., Fuchs, H., Gailus-Durner, V., Wells, S., et al. (2013). A comparative phenotypic and genomic analysis of C57BL/6J and C57BL/6N mouse strains. Genome Biology, 14, R82. 10.1186/gb-2013-14-7-r82

Sobolevsky, A. I., Koshelev, S. G., & Khodorov, B. I. (1998). Interaction of memantine and amantadine with agonist-unbound NMDA-receptor channels in acutely isolated rat hippocampal neurons. The Journal of Physiology, 512, 47–60. 10.1111/j.1469-7793.1998.047bf.x

Song, X., Jensen, M. Ø., Jogini, V., Stein, R. A., Lee, C.-H., Mchaourab, H. S., et al. (2018). Mechanism of NMDA receptor channel block by MK-801 and memantine. Nature, 556, 515–519. 10.1038/s41586-018-0039-9

Steigerwald, R., Epstein, M., Chou, T.-H., Simorowski, N., & Furukawa, H. (2026). Molecular mechanism of calcium permeability and magnesium block in NMDA receptors. Nature Neuroscience, 29, 1293–1302. 10.1038/s41593-026-02283-3

Strehlow, V., Heyne, H. O., Vlaskamp, D. R. M., Marwick, K. F. M., Rudolf, G., de Bellescize, J., et al. (2019). GRIN2A-related disorders: genotype and functional consequence predict phenotype. Brain, 142, 80–92. 10.1093/brain/awy304

Stroebel, D., Casado, M., & Paoletti, P. (2018). Triheteromeric NMDA receptors: from structure to synaptic physiology. Current Opinion in Physiology, 2, 1–12. 10.1016/j.cophys.2017.12.004

Toki, H., Ichikawa, T., Mizuno-Yasuhira, A., & Yamaguchi, J.-I. (2018). A rapid and sensitive chiral LC–MS/MS method for the determination of ketamine and norketamine in mouse plasma, brain and cerebrospinal fluid applicable to the stereoselective pharmacokinetic study of ketamine. Journal of Pharmaceutical and Biomedical Analysis, 148, 288–297. 10.1016/j.jpba.2017.09.033

Ueno, H., Suemitsu, S., Murakami, S., Kitamura, N., Wani, K., Takahashi, Y., et al. (2020). Effects of haloperidol inhalation on MK-801-and memantine-induced locomotion in mice. Libyan Journal of Medicine, 15, 1808361. 10.1080/19932820.2020.1808361

Wallach, J., Kang, H., Colestock, T., Morris, H., Bortolotto, Z. A., Collingridge, G. L., et al. (2016). Pharmacological Investigations of the Dissociative ‘Legal Highs’ Diphenidine, Methoxphenidine and Analogues. PLOS ONE, 11, e0157021. 10.1371/journal.pone.0157021

Wild, A., Akyol, E., Brothwell, S., Kimkool, P., Skepper, J., Gibb, A., et al. (2013). Memantine block depends on agonist presentation at the NMDA receptor in substantia nigra pars compacta dopamine neurones. Neuropharmacology, 73, 138–146. 10.1016/j.neuropharm.2013.05.013

Wollmuth, L. P., Kuner, T., & Sakmann, B. (1998). Adjacent asparagines in the NR2-subunit of the NMDA receptor channel control the voltage-dependent block by extracellular Mg2+. The Journal of Physiology, 506, 13–32. 10.1111/j.1469-7793.1998.013bx.x

Wu, J., Zou, H., Strong, J. A., Yu, J., Zhou, X., Xie, Q., et al. (2005). Bimodal effects of MK-801 on locomotion and stereotypy in C57BL/6 mice. Psychopharmacology, 177, 256–263. 10.1007/s00213-004-1944-1

Zhang, M., Feng, J., Xie, C., Song, N., Jin, C., Wang, J., et al. (2025). Assembly and architecture of endogenous NMDA receptors in adult cerebral cortex and hippocampus. Cell, 188, 1198– 1207.e13. 10.1016/j.cell.2025.01.004

Zhang, Y., Ye, F., Zhang, T., Lv, S., Zhou, L., Du, D., et al. (2021). Structural basis of ketamine action on human NMDA receptors. Nature, 596, 301–305. 10.1038/s41586-021-03769-9

Zhou, C., & Tajima, N. (2023). Structural insights into NMDA receptor pharmacology. Biochemical Society Transactions, 51, 1713–1731. 10.1042/bst20230122

