## Supplementary figures and images for "Dizocilpine-derived K2060 exhibits enhanced blockage at NMDA receptors with impaired Mg^2+^ block and provides strong protection against audiogenic seizures"

### Supplementary Figures

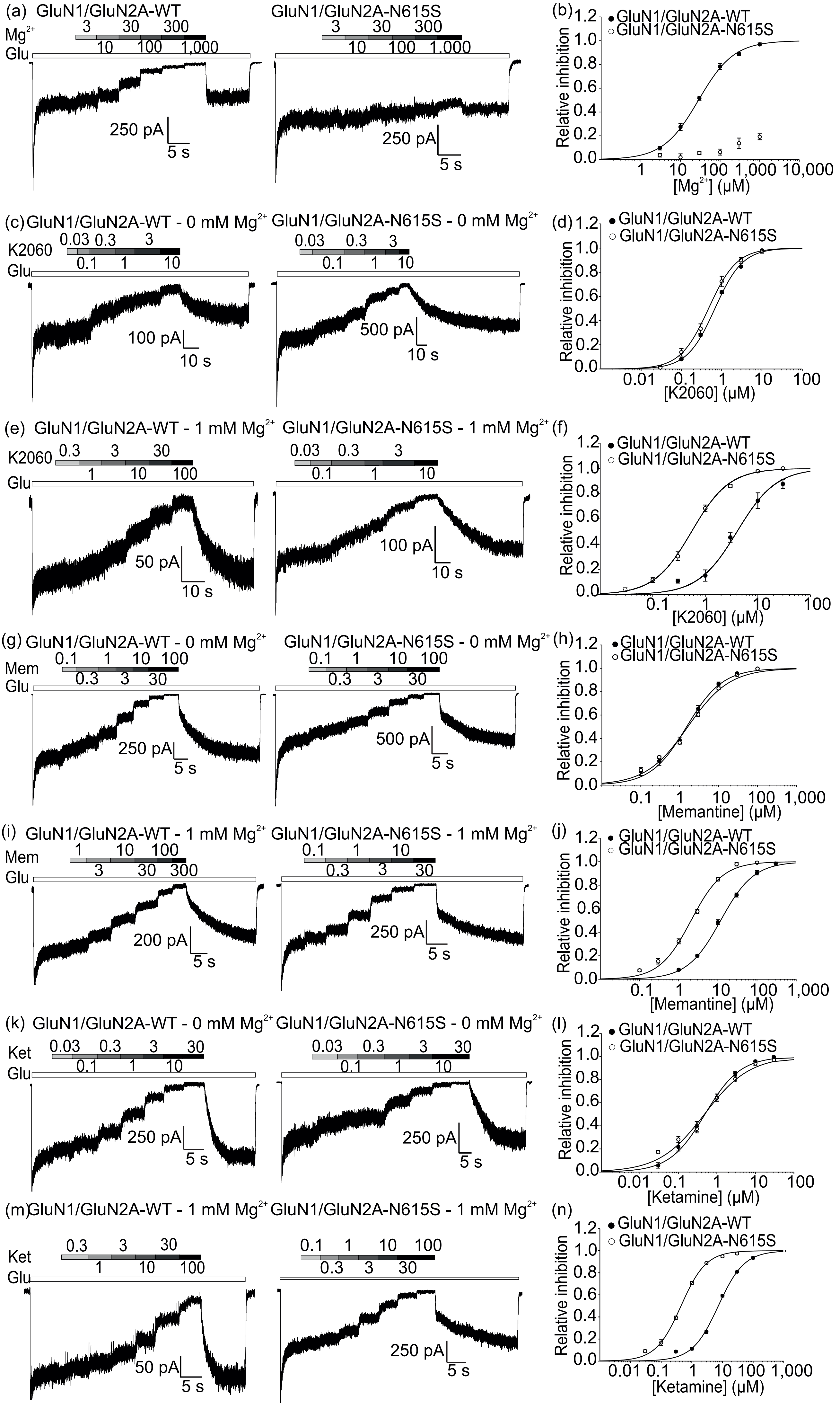

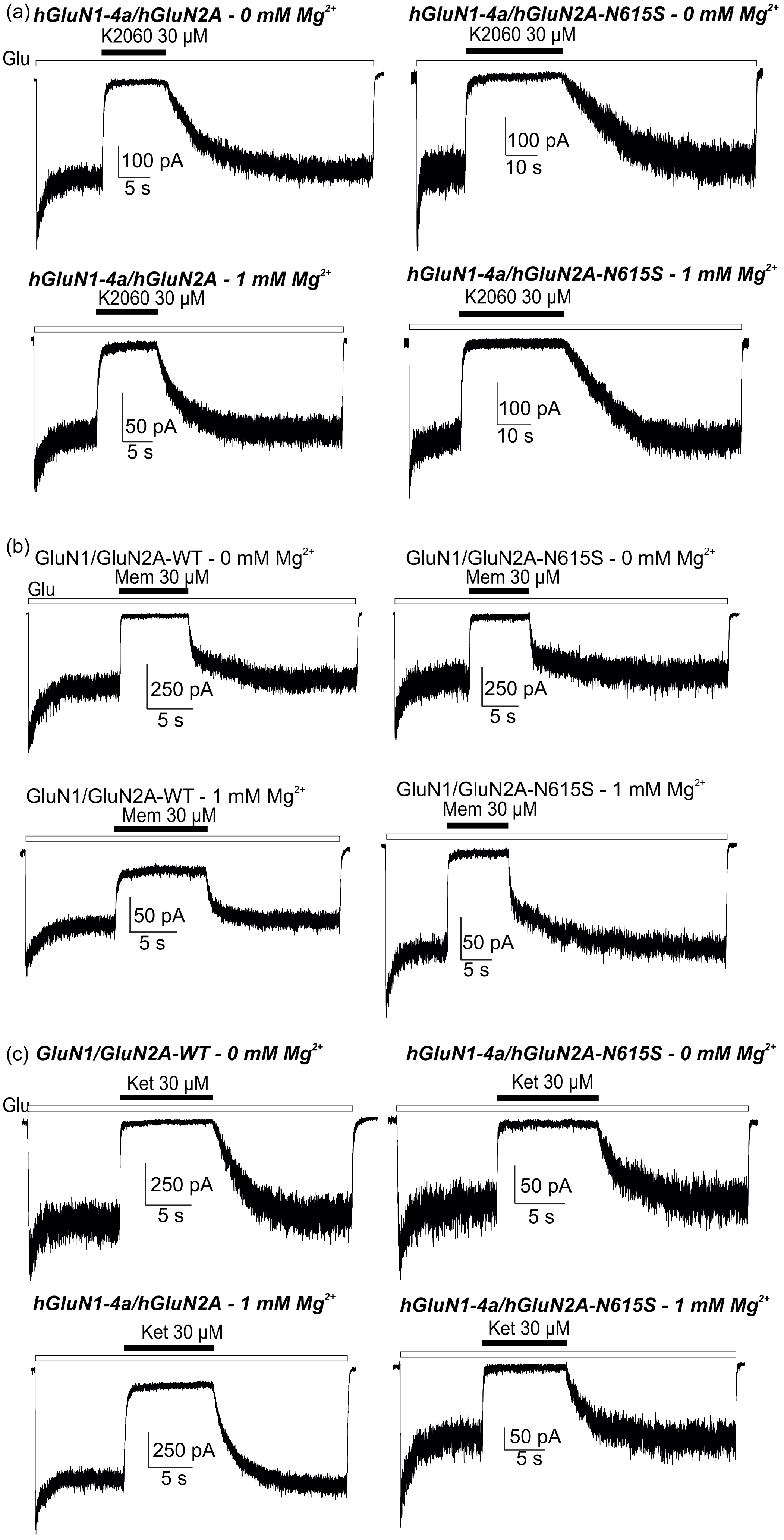
