## Supplementary Tables and supplementary figure legend for "Dizocilpine-derived K2060 exhibits enhanced blockage at NMDA receptors with impaired Mg^2+^ block and provides strong protection against audiogenic seizures"

SUPPLEMENTARY FIGURE 1 Concentration-dependent inhibition of GluN1/GluN2A-WT and GluN1/GluN2A-N615S receptors. (a, b) Representative whole-cell patch-clamp recordings and concentration-response curves for $Mg^{2+}$ inhibition of GluN1/GluN2A-WT and GluN1/GluN2A-N615S receptors expressed in HEK293T cells. (c–f) Memantine inhibition in the absence of extracellular $Mg^{2+}$ (c, d) and in the presence of 1 mM $Mg^{2+}$ (e, f). (g–j) Ketamine inhibition in the absence of extracellular $Mg^{2+}$ (g, h) and in the presence of 1 mM $Mg^{2+}$ (i, j). (k–n) K2060 inhibition in the absence of extracellular $Mg^{2+}$ (k, l) and in the presence of 1 mM $Mg^{2+}$ (m, n). Currents were evoked by 1 mM L-glutamate, as indicated by open bars, and increasing blocker concentrations are indicated by filled bars. Concentration-response data are shown as mean ± SEM. $IC_{50}$ values were determined by fitting concentration-response relationships from individual cell recordings using *Equation 1* and are reported in Table 1; full statistical details are provided in Supplementary Table S5.

SUPPLEMENTARY FIGURE 2 (a–c) Representative whole-cell current responses recorded from HEK293T cells expressing recombinant GluN1/GluN2A-WT or GluN1/GluN2A-N615S receptors. Traces show inhibition by 30 μM K2060 (a), memantine (Mem; b) or ketamine (Ket; c) during receptor activation in the absence or presence of 1 mM extracellular $Mg^{2+}$, as indicated above each trace. White bars indicate 1 mM L-glutamate application, and black bars indicate blocker application. Glycine was present as the co-agonist as described in *Methods*. Related data are shown in Figure 2.

Supplementary Table S1
Summary of the time constants of the onset and offset of inhibition by K2060 measured in HEK293T cells expressing the indicated NMDARs (related data are shown in Figure 2a–c and Figure 6c,d)

| Receptor | $Mg^{2+}$ | K2060 (μM) | $\tau_{on}$ (ms) | $\tau_{off}$ (ms) | $\tau_{fast}$ (ms) | $\tau_{slow}$ (ms) | $A_{fast}$ | $A_{slow}$ |
|---|---|---|---|---|---|---|---|---|
| GluN1/GluN2A-WT | 0 | 10 | 384.92 ± 41.35 (9) | 8,943.33 ± 1043.33 (9) | 829.47 ± 259.45 (9) | 9,723.87 ± 1062.08 (9) | 0.10 ± 0.04 (9) | 0.90 ± 0.04 (9) |
| | 1 | 10 | 1,379.12 ± 110.18 (5) | 4,494.98 ± 372.52 (5) | 806.33 ± 256.52 (5) | 5,429.33 ± 777.85 (5) | 0.18 ± 0.06 (5) | 0.82 ± 0.06 (5) |
| GluN1/GluN2A-N615S | 0 | 10 | 1,337.79 ± 104.54 (9) | 14,695.37 ± 793.90 (8) | - | - | - | - |
| | 1 | 10 | 1,535.64 ± 150.62 (9) | 15,208.5 ± 330.89 (9) | - | - | - | - |
| GluN1/GluN2A-WT | 0 | 30 | 233.53 ± 21.27 (8) | 8,718.21 ± 1182.42 (8) | 833.28 ± 387.19 (8) | 8,718.21 ± 1182.42 (8) | 0.04 ± 0.03 (8) | 0.96 ± 0.03 (8) |
| | 1 | 30 | 608.40 ± 29.42 (5) | 4,396.06 ± 500.64 (5) | 845.05 ± 257.65 (5) | 5,347.77 ± 511.65 (5) | 0.22 ± 0.03 (5) | 0.78 ± 0.03 (5) |
| GluN1/GluN2A-N615S | 0 | 30 | 599.89 ± 36.95 (12) | 16,222.61 ± 531.32 (8) | - | - | - | - |
| | 1 | 30 | 521.48 ± 24.70 (11) | 15,703.40 ± 1192.75 (7) | - | - | - | - |
| GluN1/GluN2A-N615K | 0 | 30 | 544.84 ± 49.09 (5) | 19,188.78 ± 762.19 (4) | - | - | - | - |
| | 1 | 30 | 429.92 ± 27.49 (5) | 18,411.30 ± 1,525.96 (4) | - | - | - | - |
| GluN1/GluN2A-WT | 0 | 100 | 98.35 ± 10.31 (10) | 9,238.19 ± 1230.01 (10) | 1,632.14 ± 432.60 (10) | 9,744.04 ± 1273.23 (10) | 0.04 ± 0.06 (10) | 0.96 ± 0.06 (10) |
| | 1 | 100 | 291.92 ± 48.91 (5) | 4,866.35 ± 256.85 (4) | 912.15 ±291.75 (4) | 5,979.67 ± 733.15 (4) | 0.19 ± 0.13 (4) | 0.81 ± 0.13 (4) |

| | | | | | | | | |
|---|---|---|---|---|---|---|---|---|
| GluN1/GluN2A-N615S | 0 | 100 | 275.24 ± 26.34 (12) | 15,103.57 ± 1388.96 (12) | - | - | - | - |
| | 1 | 100 | 263.09 ± 28.66 (10) | 14,115.80 ± 671.03 (8) | - | - | - | - |

Time constants for onset ($\tau_{on}$) and offset ($\tau_{off}$) of inhibition by K2060 were obtained from whole-cell L-glutamate-evoked current responses in HEK293T cells expressing the indicated receptors in external solutions containing 0 $Mg^{2+}$ or 1 mM $Mg^{2+}$ and the K2060 concentrations shown. $\tau_{off}$ was fitted with a double-exponential function, yielding fast and slow components ($\tau_{fast}$, $\tau_{slow}$) and their relative amplitudes ($A_{fast}$, $A_{slow}$); $\tau_{off}$ values are the corresponding weighted time constants ($\tau_w$). Data are presented as mean ± SEM; *n* in parentheses indicates the number of recorded cells.

Supplementary Table S2

Summary of the time constants of the onset and offset of inhibition by memantine measured in HEK293T cells expressing the indicated NMDARs (related data are shown in Figure 2d–f and Figure 6c,d.)

| Receptor | $Mg^{2+}$ | Memantine (μM) | $\tau_{on}$ (ms) | $\tau_{off}$ (ms) | $\tau_{fast}$ (ms) | $\tau_{slow}$ (ms) | $A_{fast}$ | $A_{slow}$ |
|---|---|---|---|---|---|---|---|---|
| GluN1/GluN2A-WT | 0 | 10 | 102.34 ± 11.74 (6) | 959.02 ± 18.42 (5) | 228.85 ± 42.56 (5) | 1,691.77 ± 137.13 (5) | 0.49 ± 0.05 (5) | 0.51 ± 0.05 (5) |
| | 1 | 10 | 287.57 ± 41.86 (5) | 632.13 ± 50.73 (5) | 136.52 ± 36.60 (5) | 816.15 ± 112.86 (5) | 0.38 ± 0.08 (5) | 0.62 ± 0.08 (5) |
| GluN1/GluN2A-N615S | 0 | 10 | 101.35 ± 16.35 (6) | 971.90 ± 120.52 (7) | 214.95 ± 18.75 (7) | 2,383.51 ± 388.78 (7) | 0.63 ± 0.05 (7) | 0.37 ± 0.05 (7) |
| | 1 | 10 | 178.01 ± 24.37 (6) | 1,564.16 ± 189.58 (5) | 218.23 ± 43.67 (5) | 2,966.12 ± 472.90 (5) | 0.55 ± 0.06 (5) | 0.45 ± 0.06 (5) |
| GluN1/GluN2A-WT | 0 | 30 | 52.52 ± 5.24 (6) | 2,096.67 ± 192.21 (5) | 338.36 ± 63.30 (5) | 4,381.31 ± 301.19 (5) | 0.56 ± 0.06 (5) | 0.44 ± 0.06 (5) |
| | 1 | 30 | 160.43 ± 13.78 (6) | 616.37 ± 106.18 (6) | 152.53 ± 45.23 (6) | 1,005.80 ± 209.93 (6) | 0.44 ± 0.06 (6) | 0.56 ± 0.06 (6) |
| GluN1/GluN2A-N615S | 0 | 30 | 53.95 ± 4.25 (7) | 2,199.55 ± 1155.06 (7) | 236.59 ± 22.41 (7) | 5,404.26 ± 370.71 (7) | 0.61 ± 0.04 (7) | 0.39 ± 0.04 (7) |
| | 1 | 30 | 80.96 ± 4.03 (6) | 1,777.11 ± 189.20 (4) | 178.74 ± 14.81 (4) | 3,668.80 ± 540.36 (4) | 0.53 ± 0.04 (4) | 0.47 ± 0.04 (4) |
| GluN1/GluN2A-N615K | 0 | 30 | 555.55 ± 65.94 (6) | 1,791.52 ± 235.47 (6) | 342.08 ± 60.18 (6) | 2,966.12 ± 490.02 (6) | 0.43 ± 0.06 (6) | 0.57 ± 0.06 (6) |
| | 1 | 30 | 671.54 ± 26.88 (6) | 2064.92 ± 278.15 (7) | 345.08 ± 32.07 (7) | 3,467.87 ± 521.83 (7) | 0.37 ± 0.03 (7) | 0.63 ± 0.03 (7) |
| GluN1/GluN2A-WT | 0 | 100 | 27.06 ± 1.61 (5) | 3,772.32 ± 775.38 (4) | 293.11 ± 60.74 (4) | 5,740.01 ± 798.88 (4) | 0.39 ± 0.07 (4) | 0.61 ± 0.07 (4) |
| | 1 | 100 | 86.93 ± 8.41 (5) | 908.44 ± 134.23 (5) | 200.16 ± 31.64 (5) | 1,789.79 ± 211.72 (5) | 0.55 ± 0.06 (5) | 0.45 ± 0.06 (5) |

| | | | | | | | | |
|---|---|---|---|---|---|---|---|---|
| GluN1/GluN2A-N615S | 0 | 100 | 26.48 ± 2.77 (7) | 2,807.46 ± 206.63 (7) | 225.35 ± 26.86 (7) | 4,896.13 ± 26.86 (7) | 0.44 ± 0.03 (7) | 0.56 ± 0.03 (7) |
| | 1 | 100 | 42.18 ± 7.46 (5) | 2,400.87 ± 242.40 (6) | 136.31 ± 13.46 (6) | 3,842.20 ± 381.78 (6) | 0.39 ± 0.02 (6) | 0.61 ± 0.02(6) |

Time constants for onset ($\tau_{on}$) and offset ($\tau_{off}$) of inhibition by memantine were obtained from whole-cell glutamate-evoked currents in HEK293T cells expressing the indicated receptors in external solutions containing 0 $Mg^{2+}$ or 1 mM $Mg^{2+}$ and the memantine concentrations shown. $\tau_{off}$ was fitted with a double-exponential function, yielding fast and slow components ($\tau_{fast}$, $\tau_{slow}$) and their relative amplitudes ($A_{fast}$, $A_{slow}$); $\tau_{off}$ values are the corresponding weighted time constants ($\tau_w$). Data are presented as mean ± SEM; *n* in parentheses indicates the number of recorded cells.

Supplementary Table S3

Summary of the time constants of the onset and offset of inhibition by ketamine measured in HEK293T cells expressing the indicated NMDARs (related data are shown in Figure 2g–i and Figure 6c,d.)

| Receptor | $Mg^{2+}$ | Ketamine (μM) | $\tau_{on}$ (ms) | $\tau_{off}$ (ms) | $\tau_{fast}$ (ms) | $\tau_{slow}$ (ms) | $A_{fast}$ | $A_{slow}$ |
|---|---|---|---|---|---|---|---|---|
| GluN1/GluN2A-WT | 0 | 10 | 167.99 ± 8.62 (6) | 1,997.32 ± 272.872(6) | 332.06 ± 54.21 (6) | 2,602.54 ± 404.47 (6) | 0.26 ± 0.05 (6) | 0.74 ± 0.05 (6) |
| | 1 | 10 | 531.39 ± 59.44 (7) | 2,090.83 ± 273.96 (6) | 395.64 ± 41.65 6) | 2,543.71 ± 317.01 (6) | 0.19 ± 0.08 (6) | 0.81 ± 0.08 (6) |
| GluN1/GluN2A-N615S | 0 | 10 | 178.50 ± 30.39 (8) | 2,456.05 ± 214.14 (6) | 253.83 ± 51.14 (6) | 2,819.22 ± 296.60 (6) | 0.13 ± 0.03 (6) | 0.87 ± 0.03 (6) |
| | 1 | 10 | 234.26 ± 20.10 (9) | 2,685.34 ± 572.46 (5) | 261.27 ± 115.41 (5) | 3,056.44 ± 581.64 (5) | 0.17 ± 0.07 (5) | 0.83 ± 0.07 (5) |
| GluN1/GluN2A-WT | 0 | 30 | 135.46 ± 16.77 (10) | 2,156.58 ± 210.49 (6) | 349.51 ± 98.66 (6) | 2,375.52 ± 162.18 (6) | 0.12 ± 0.05 (6) | 0.88 ± 0.05 (6) |
| | 1 | 30 | 370.41 ± 65.19 (6) | 1,944.75 ± 200.20 (6) | 459.86 ± 93.28 (6) | 2,442.47 ± 303.60 (6) | 0.25 ± 0.05 (6) | 0.75 ± 0.05 (6) |
| GluN1/GluN2A-N615S | 0 | 30 | 134.31 ± 22.21 (8) | 2,607.52 ± 247.01 (8) | 225.09 ± 65.26 (8) | 2,862.50 ± 272.62 (9) | 0.10 ± 0.01 (8) | 0.90 ± 0.01 (8) |
| | 1 | 30 | 113.30 ±14.32 (5) | 2,656.17 ± 605.74 (4) | 268.50 ± 104.68 (4) | 2,946.75 ± 634.69 (4) | 0.12 ± 0.04 (4) | 0.88 ± 0.04 (4) |
| GluN1/GluN2A-N615K | 0 | 30 | 457.64 ± 47.60 (6) | 9,351.66 ± 885.49 (6) | - | - | - | - |
| | 1 | 30 | 419.12 ± 95.15 (4) | 8,660.09 ± 1,613.74 (4) | - | - | - | - |
| GluN1/GluN2A-WT | 0 | 100 | 67.46 ± 10.56 (12) | 2,490.43 ± 394.56 (7) | 376.64 ± 118.14 (7) | 2,751.83 ± 414.35 (7) | 0.12 ± 0.09 (5) | 0.88 ± 0.09 (5) |
| | 1 | 100 | 223.30 ± 28.99 (7) | 2,394.49 ± 210.19 (5) | 519.56 ± 87.39 (5) | 2,866.33 ± 135.58 (5) | 0.21 ± 0.04 (5) | 0.79 ± 0.04 (5) |
| GluN1/GluN2A-N615S | 0 | 100 | 59.69 ± 13.85 (8) | 2,474.61 ± 362.92 (5) | 453.02 ± 91.61 (5) | 2,846.21 ± 261.96 (5) | 0.28 ± 0.08 (5) | 0.82 ± 0.08 (5) |
| | 1 | 100 | 59.27 ± 7.82 (5) | 2,922.11 ± 445.81 (4) | 512.69 ± 237.33 (4) | 3,872.73 ± 588.18 (4) | 0.25 ± 0.16 (4) | 0.79 ± 0.04 (4) |

Time constants for onset ($\tau_{on}$) and offset ($\tau_{off}$) of inhibition by ketamine were obtained from whole-cell L-glutamate-evoked current responses in HEK293T cells expressing the indicated receptors in external solutions containing 0 $Mg^{2+}$ or 1 mM $Mg^{2+}$ and the ketamine concentrations shown. $\tau_{off}$ was fitted with a double-exponential function, yielding fast and slow components ($\tau_{fast}$, $\tau_{slow}$) and their relative amplitudes ($A_{fast}$, $A_{slow}$); $\tau_{off}$ values are the corresponding weighted time constants ($\tau_w$). Data are presented as mean ± SEM; *n* in parentheses indicates the number of recorded cells.

Supplementary Table S4

Summary of the time constants of the offset kinetic from memantine, ketamine and K2060 inhibition measured with the double-pulse protocol (related data are shown in Figure 3)

| Receptor | Antagonist | $Mg^{2+}$ (mM) | $\tau_w$ | $\tau_{fast}$ | $\tau_{slow}$ | $A_{fast}$ | $A_{slow}$ | n |
|---|---|---|---|---|---|---|---|---|
| GluN1/GluN2A-WT | K2060 | 0 | 7,761.25± 243.26 | 284.70 ± 81.06 | 10,063.48± 464.22 | 0.23±0.05 | 0.77±0.05 | 4 |
| | Mem | 0 | 1,301.56 ±195.21 | 255.51 ±50.82 | 2,052.76± 311.24 | 0.42 ± 0.02 | 0.58 ± 0.02 | 5 |
| | Ket | 0 | 6,197.27± 921.78 | 255.75 ±61.34 | 8,290.12± 1,170.75 | 0.26±0.02 | 0.74±0.02 | 5 |
| GluN1/GluN2A-N615S | K2060 | 0 | 14,382.71± 1,951.17 | 194.94 ±13.63 | 17,082.01± 2,509.34 | 0.15±0.03 | 0.85±0.03 | 4 |
| | Mem | 0 | 1,372.58±129.39 | 194.13 ±34.27 | 2,780.42±157.03 | 0.53±0.06 | 0.47±.06 | 5 |
| | Ket | 0 | 5,798.49± 503.45 | 265.62 ±93.06 | 7,553.29± 650.64 | 0.24±0.03 | 0.76±0.03 | 5 |
| GluN1/GluN2A-WT | K2060 | 1 | 6,097.52± 677.42 | 280.33 ±124.21 | 8,706.45± 1,202.93 | 0.30±0.03 | 0.70±0.03 | 4 |
| | Mem | 1 | 2,687.22±600.12 | 240.61 ±61.98 | 4,291.09±944.11 | 0.40±0.01 | 0.60±0.01 | 5 |
| | Ket | 1 | 5,292.91±521.86 | 367.25 ±200.93 | 7,179.29±856.49 | 0.27±0.02 | 0.73±0.02 | 4 |
| GluN1/GluN2A-N615S | K2060 | 1 | 13,900.09± 1,158.69 | 176.64 ±13.55 | 16,988.61± 1,685.31 | 0.18±0.02 | 0.82±0.02 | 4 |
| | Mem | 1 | 1,429.82± 205.52 | 173.50 ±44.89 | 2,649.75± 270.60 | 0.49±0.07 | 0.51±0.07 | 6 |
| | Ket | 1 | 7,249.54± 655.06 | 323.46 ± 60.61 | 8,818.37±703.20 | 0.19±0.02 | 0.81±0.02 | 4 |

Weighted recovery time constants ($\tau_w$), fast and slow components ($\tau_{off}$-fast, $\tau_{off}$-slow) and their relative amplitudes ($A_{fast}$, $A_{slow}$) were obtained by fitting double-exponential functions to NMDA-evoked current responses during the second pulse of the double-pulse protocol. Recordings were made from HEK293T cells expressing GluN1/GluN2A or GluN1/GluN2A-N615S receptors in external solutions containing 0 or 1 mM $Mg^{2+}$, as indicated. $A_{slow}$ reflects the trapped fraction of block persisting after the agonist-free wash. Data are mean ± SEM; *n* denotes the number of cells.

Supplementary Table S5

Statistical analysis supporting Table 1

| Compound | Comparison | Test | Statistic | *P* value |
|---|---|---|---|---|
| Memantine | Receptor variant | Two-way ANOVA | F(2,34) = 154.205 | <0.001 |
| Memantine | $Mg^{2+}$ condition | Two-way ANOVA | F(1,34) = 170.423 | <0.001 |
| Memantine | Receptor × $Mg^{2+}$ | Two-way ANOVA | F(2,34) = 128.188 | <0.001 |
| Memantine | WT, 0 vs 1 mM $Mg^{2+}$ | Holm-Sidak | t(34) = 20.361 | <0.001 |
| Memantine | N615S, 0 vs 1 mM $Mg^{2+}$ | Holm-Sidak | t(34) = 0.972 | 0.338 |
| Memantine | N615K, 0 vs 1 mM $Mg^{2+}$ | Holm-Sidak | t(34) = 0.938 | 0.355 |
| Memantine | WT vs N615S at 0 mM $Mg^{2+}$ | Holm-Sidak | t(34) = 0.203 | 0.841 |
| Memantine | WT vs N615K at 0 mM $Mg^{2+}$ | Holm-Sidak | t(34) = 12.332 | <0.001 |
| Memantine | WT vs N615S at 1 mM $Mg^{2+}$ | Holm-Sidak | t(34) = 19.186 | <0.001 |
| Memantine | WT vs N615K at 1 mM $Mg^{2+}$ | Holm-Sidak | t(34) = 7.789 | <0.001 |
| Ketamine | WT, 0 vs 1 mM $Mg^{2+}$ | Welch t + Holm | t(4.623) = -29.296 | <0.001 |
| Ketamine | N615S, 0 vs 1 mM $Mg^{2+}$ | Welch t + Holm | t(8.074) = 0.455 | 1.000 |
| Ketamine | N615K, 0 vs 1 mM $Mg^{2+}$ | Welch t + Holm | t(7.083) = -0.246 | 1.000 |

| | | | | |
|---|---|---|---|---|
| Ketamine | WT vs N615S at 0 mM $Mg^{2+}$ | Welch t + Holm | t(9.703) = 0.171 | 1.000 |
| Ketamine | WT vs N615K at 0 mM $Mg^{2+}$ | Welch t + Holm | t(6.191) = -0.730 | 1.000 |
| Ketamine | WT vs N615S at 1 mM $Mg^{2+}$ | Welch t + Holm | t(4.117) = 30.412 | <0.001 |
| Ketamine | WT vs N615K at 1 mM $Mg^{2+}$ | Welch t + Holm | t(4.974) = 28.153 | <0.001 |
| K2060 | WT, 0 vs 1 mM $Mg^{2+}$ | Mann-Whitney + Holm | U = 0.000 | 0.040 |
| K2060 | N615S, 0 vs 1 mM $Mg^{2+}$ | Mann-Whitney + Holm | U = 30.000 | 0.452 |
| K2060 | N615K, 0 vs 1 mM $Mg^{2+}$ | Mann-Whitney + Holm | U = 10.000 | 0.452 |
| K2060 | WT vs N615S at 0 mM $Mg^{2+}$ | Mann-Whitney + Holm | U = 10.000 | 0.448 |
| K2060 | WT vs N615K at 0 mM $Mg^{2+}$ | Mann-Whitney + Holm | U = 5.000 | 0.452 |
| K2060 | WT vs N615S at 1 mM $Mg^{2+}$ | Mann-Whitney + Holm | U = 0.000 | 0.003 |
| K2060 | WT vs N615K at 1 mM $Mg^{2+}$ | Mann-Whitney + Holm | U = 0.000 | 0.026 |

Statistical comparisons were performed on fitted $IC_{50}$ values from individual recorded cells. For memantine, normality and equal variance assumptions were met, and data were analysed using two-way ANOVA with receptor variant and $Mg^{2+}$ condition as factors, followed by Holm–Sidak *post hoc* tests. For ketamine, normality was met, but equal variance was not; therefore, predefined pairwise comparisons were performed using Welch's *t*-tests with Holm correction. For K2060, normality was not met; therefore, predefined pairwise comparisons were performed using Mann–Whitney rank-sum tests with Holm correction.

Supplementary Table S6
Statistical analysis of kinetic data (related data are shown in Figure 2)

| Figure | Compound/concentration | Comparison | Test | *P* value |
|---|---|---|---|---|
| Fig. 2b | K2060; $\tau_{on}$; 10 µM | $Mg^{2+}$ effect in WT | unpaired Student's t-test | <0.001 |
| Fig. 2b | K2060; $\tau_{on}$; 10 µM | WT vs N615S at 0 mM $Mg^{2+}$ | Welch's t-test | <0.001 |
| Fig. 2b | K2060; $\tau_{on}$; 30 µM | $Mg^{2+}$ effect in WT | unpaired Student's t-test | <0.001 |
| Fig. 2b | K2060; $\tau_{on}$; 30 µM | $Mg^{2+}$ effect in N615S | unpaired Student's t-test | 0.039 |
| Fig. 2b | K2060; $\tau_{on}$; 30 µM | WT vs N615S at 0 mM $Mg^{2+}$ | unpaired Student's *t*-test | <0.001 |
| Fig. 2b | K2060; $\tau_{on}$; 30 µM | WT vs N615S at 1 mM $Mg^{2+}$ | unpaired Student's *t*-test | 0.039 |
| Fig. 2b | K2060; $\tau_{on}$; 100 µM | $Mg^{2+}$ effect in WT | Welch's *t*-test | 0.047 |
| Fig. 2b | K2060; $\tau_{on}$; 100 µM | WT vs N615S at 0 mM $Mg^{2+}$ | Welch's *t*-test | <0.001 |
| Fig. 2c | K2060; $\tau_{off}$; 10 µM | $Mg^{2+}$ effect in WT | unpaired Student's *t*-test | 0.024 |
| Fig. 2c | K2060; $\tau_{off}$; 10 µM | WT vs N615S at 0 mM $Mg^{2+}$ | unpaired Student's *t*-test | 0.002 |
| Fig. 2c | K2060; $\tau_{off}$; 10 µM | WT vs N615S at 1 mM $Mg^{2+}$ | unpaired Student's *t*-test | <0.001 |
| Fig. 2c | K2060; $\tau_{off}$; 30 µM | Mg2+ effect in WT | Mann–Whitney rank-sum test | 0.003 |
| Fig. 2c | K2060; $\tau_{off}$; 30 µM | WT vs N615S at 0 mM $Mg^{2+}$ | Mann–Whitney rank-sum test | <0.001 |
| Fig. 2c | K2060; $\tau_{off}$; 30 µM | WT vs N615S at 1 mM $Mg^{2+}$ | unpaired Student's *t*-test | <0.001 |
| Fig. 2c | K2060; $\tau_{off}$; 100 µM | $Mg^{2+}$ effect in WT | Mann–Whitney rank-sum test | 0.012 |
| Fig. 2c | K2060; $\tau_{off}$; 100 µM | WT vs N615S at 0 mM $Mg^{2+}$ | Mann–Whitney rank-sum test | 0.004 |
| Fig. 2c | K2060; $\tau_{off}$; 100 µM | WT vs N615S at 1 mM $Mg^{2+}$ | Mann–Whitney rank-sum test | 0.012 |
| Fig. 2e | Memantine; $\tau_{on}$; 10 µM | $Mg^{2+}$ effect in WT | unpaired Student's t-test | 0.005 |
| Fig. 2e | Memantine; $\tau_{on}$; 30 µM | $Mg^{2+}$ effect in WT | Welch's t-test | <0.001 |
| Fig. 2e | Memantine; $\tau_{on}$; 30 µM | $Mg^{2+}$ effect in N615S | unpaired Student's t-test | 0.002 |
| Fig. 2e | Memantine; $\tau_{on}$; 30 µM | WT vs N615S at 1 mM $Mg^{2+}$ | Welch's t-test | 0.003 |
| Fig. 2e | Memantine; $\tau_{on}$; 100 µM | $Mg^{2+}$ effect in WT | unpaired Student's t-test | <0.001 |
| Fig. 2e | Memantine; $\tau_{on}$; 100 µM | WT vs N615S at 1 mM $Mg^{2+}$ | unpaired Student's t-test | 0.009 |
| Fig. 2f | Memantine; $\tau_{off}$; 10 µM | $Mg^{2+}$ effect in WT | unpaired Student's t-test | 0.001 |
| Fig. 2f | Memantine; $\tau_{off}$; 10 µM | WT vs N615S at 1 mM $Mg^{2+}$ | Welch's t-test | 0.037 |
| Fig. 2f | Memantine; $\tau_{off}$; 30 µM | $Mg^{2+}$ effect in WT | unpaired Student's t-test | <0.001 |
| Fig. 2f | Memantine; $\tau_{off}$; 30 µM | WT vs N615S at 1 mM $Mg^{2+}$ | unpaired Student's t-test | 0.001 |
| Fig. 2f | Memantine; $\tau_{off}$; 100 µM | WT vs N615S at 1 mM $Mg^{2+}$ | unpaired Student's t-test | 0.003 |
| Fig. 2h | Ketamine; $\tau_{on}$; 10 µM | $Mg^{2+}$ effect in WT | Welch's t-test | 0.005 |
| Fig. 2h | Ketamine; $\tau_{on}$; 10 µM | WT vs N615S at 1 mM $Mg^{2+}$ | unpaired Student's t-test | 0.016 |
| Fig. 2h | Ketamine; $\tau_{on}$; 30 µM | $Mg^{2+}$ effect in WT | unpaired Student's t-test | 0.003 |
| Fig. 2h | Ketamine; $\tau_{on}$; 30 µM | WT vs N615S at 1 mM $Mg^{2+}$ | unpaired Student's t-test | 0.011 |
| Fig. 2h | Ketamine; $\tau_{on}$; 100 µM | $Mg^{2+}$ effect in WT | Mann–Whitney rank-sum test | <0.001 |
| Fig. 2h | Ketamine; $\tau_{on}$; 100 µM | WT vs N615S at 1 mM $Mg^{2+}$ | unpaired Student's t-test | <0.001 |

Statistical analysis of onset and recovery kinetics for memantine, ketamine and K2060 inhibition at GluN1/GluN2A-WT and GluN1/GluN2A-N615S receptors. The table includes only significant predefined comparisons corresponding to the statistical annotations shown in Figure 1. Comparisons were performed on $\tau_{on}$ and $\tau_{off}$ values from individual recorded cells. Tests were selected according to normality and equality of variance, as described in Methods.

Supplementary Table S7
Statistical analysis of trapping analysis (related data are shown in Figure 3)

| Analysis | Groups tested | Test | Statistic (df) | *P* value |
|---|---|---|---|---|
| Compound effect in WT, 0 mM $Mg^{2+}$ | Mem *n* = 5; Ket *n* = 5; K2060 *n* = 4 | One-way ANOVA + Holm-Sidak | F(2,11)=11.479 | 0.002 |
| Compound effect in N615S, 0 mM $Mg^{2+}$ | Mem *n* = 5; Ket *n* = 5; K2060 *n* = 4 | One-way ANOVA + Holm-Sidak | F(2,11)=16.996 | <0.001 |
| Compound effect in WT, 1 mM $Mg^{2+}$ | Mem *n* = 5; Ket *n* = 4; K2060 *n* = 4 | One-way ANOVA + Holm-Sidak | F(2,10)=12.90 | 0.002 |
| Compound effect in N615S, 1 mM $Mg^{2+}$ | Mem *n* = 6; Ket *n* = 4; K2060 *n* = 4 | One-way ANOVA + Holm-Sidak | F(2,11)=11.43 | 0.002 |
| Receptor/$Mg^{2+}$ effect on Mem trapping | WT 0, WT 1, N615S 0, N615S 1 mM $Mg^{2+}$ | Mann-Whitney tests + Holm correction | - | >0.05 |
| Receptor/$Mg^{2+}$ effect on Ket trapping | WT 0, WT 1, N615S 0, N615S 1 mM $Mg^{2+}$ | Two-way ANOVA | - | Receptor: *P*=0.083; $Mg^{2+}$: *P*=0.409; interaction: *P*=0.313 |
| Receptor/$Mg^{2+}$ effect on K2060 trapping | WT 0, WT 1, N615S 0, N615S 1 mM $Mg^{2+}$ | Mann-Whitney tests + Holm correction | - | >0.05 |

Supplementary Table S8
Statistical analysis of time-course inhibition data and final 5-min inhibition in acute hippocampal slices

| Figure | Endpoint / dataset | n values | Comparison or effect | Test | Statistic (df) | *P* value |
|---|---|---|---|---|---|---|
| Fig. 4a | Baseline-normalized inhibition, K2060 | WT *n* = 6 slices; N615S *n* = 6 slices | Genotype effect | Two-way repeated-measures ANOVA | F(1,5) = 12.827 | 0.016 |
| Fig. 4a | Baseline-normalized inhibition, K2060 | WT *n* = 6 slices; N615S *n* = 6 slices | Genotype × time interaction | Two-way repeated-measures ANOVA | F(60,298) = 4.361 | <0.001 |
| Fig. 4b | Scaled inhibition onset, K2060 | WT *n* = 6 slices; N615S *n* = 6 slices | Genotype effect | Two-way repeated-measures ANOVA | F(1,5) = 4.041 | 0.101 |
| Fig. 4b | Scaled inhibition onset, K2060 | WT *n* = 6 slices; N615S *n* = 6 slices | Genotype × time interaction | Two-way repeated-measures ANOVA | F(60,298) = 1.958 | <0.001 |
| Fig. 4c | Baseline-normalized inhibition, memantine | WT *n* = 6 slices; N615S *n* = 6 slices | Genotype effect | Two-way repeated-measures ANOVA | F(1,10) = 5.780 | 0.037 |
| Fig. 4c | Baseline-normalized inhibition, memantine | WT *n* = 6 slices; N615S *n* = 6 slices | Genotype × time interaction | Two-way repeated-measures ANOVA | F(35,350) = 2.962 | <0.001 |
| Fig. 4d | Scaled inhibition onset, memantine | WT *n* = 6 slices; N615S *n* = 6 slices | Genotype effect | Two-way repeated-measures ANOVA | F(1,10) = 0.315 | 0.587 |
| Fig. 4d | Scaled inhibition onset, memantine | WT *n* = 6 slices; N615S *n* = 6 slices | Genotype × time interaction | Two-way repeated-measures ANOVA | F(35,350) = 0.589 | 0.971 |

| | | | | | | |
|---|---|---|---|---|---|---|
| Fig. 4e | Baseline-normalized inhibition, ketamine | WT *n* = 6 slices; N615S *n* = 6 slices | Genotype effect | Two-way repeated-measures ANOVA | F(1,5) = 53.420 | <0.001 |
| Fig. 4e | Baseline-normalized inhibition, ketamine | WT *n* = 6 slices; N615S *n* = 6 slices | Genotype × time interaction | Two-way repeated-measures ANOVA | F(45,221) = 8.875 | <0.001 |
| Fig. 4f | Scaled inhibition onset, ketamine | WT *n* = 6 slices; N615S *n* = 6 slices | Genotype effect | Two-way repeated-measures ANOVA | F(1,5) = 12.625 | 0.016 |
| Fig. 4f | Scaled inhibition onset, ketamine | WT *n* = 6 slices; N615S *n* = 6 slices | Genotype × time interaction | Two-way repeated-measures ANOVA | F(45,221) = 5.405 | <0.001 |
| Fig. 4g | Final 5-min inhibition in acute slices | n = 6 slices per drug/genotype group; total *n* = 36 slices | Genotype effect | Two-way ANOVA | F(1,30) = 41.837 | <0.001 |
| Fig. 4g | Final 5-min inhibition in acute slices | n = 6 slices per drug/genotype group; total *n* = 36 slices | Drug effect | Two-way ANOVA | F(2,30) = 42.671 | <0.001 |
| Fig. 4g | Final 5-min inhibition in acute slices | n = 6 slices per drug/genotype group; total *n* = 36 slices | Genotype × drug interaction | Two-way ANOVA | F(2,30) = 0.044 | 0.957 |
| Fig. 4g | Final 5-min inhibition in acute slices | WT *n* = 6 slices; N615S *n* = 6 slices | WT vs N615S, memantine | Holm–Sidak post hoc test | t = 3.769 | <0.001 |
| Fig. 4g | Final 5-min inhibition in acute slices | WT *n* = 6 slices; N615S *n* = 6 slices | WT vs N615S, ketamine | Holm–Sidak post hoc test | t = 3.926 | <0.001 |
| Fig. 4g | Final 5-min inhibition in acute slices | WT *n* = 6 slices; N615S *n* = 6 slices | WT vs N615S, K2060 | Holm–Sidak post hoc test | t = 3.509 | 0.001 |
| Fig. 4g | Final 5-min inhibition in acute slices | WT: memantine *n* = 6 slices; K2060 *n* = 6 slices | Memantine vs K2060 within WT | Holm–Sidak post hoc test | t = 6.085 | <0.001 |
| Fig. 4g | Final 5-min inhibition in acute slices | WT: memantine *n* = 6 slices; ketamine *n* = 6 slices | Memantine vs ketamine within WT | Holm–Sidak post hoc test | t = 5.225 | <0.001 |
| Fig. 4g | Final 5-min inhibition in acute slices | WT: ketamine *n* = 6 slices; K2060 *n* = 6 slices | Ketamine vs K2060 within WT | Holm–Sidak post hoc test | t = 0.860 | 0.397 |
| Fig. 4g | Final 5-min inhibition in acute slices | N615S: memantine *n* = 6 slices; K2060 *n* = 6 slices | Memantine vs K2060 within N615S | Holm–Sidak post hoc test | t = 5.824 | <0.001 |
| Fig. 4g | Final 5-min inhibition in acute slices | N615S: memantine *n* = 6 slices; ketamine *n* = 6 slices | Memantine vs ketamine within N615S | Holm–Sidak post hoc test | t = 5.382 | <0.001 |

| | | | | | | |
|---|---|---|---|---|---|---|
| Fig. 4g | Final 5-min inhibition in acute slices | N615S: ketamine *n* = 6 slices; K2060 *n* = 6 slices | Ketamine vs K2060 within N615S | Holm–Sidak post hoc test | t = 0.443 | 0.661 |

Time-course data were analysed separately for each blocker and normalisation approach. The main effect of time is not shown because it reflects the expected change during blocker wash-in. Baseline-normalised responses were used to compare the overall development of inhibition, whereas scaled responses were used to compare the apparent onset profile independently of final inhibition magnitude. Final 5-min inhibition was analysed by two-way ANOVA with genotype and drug as factors, followed by Holm–Sidak *post hoc* tests.

Supplementary Table S9
Statistical analysis of relative current responses obtained from HEK293T cells expressing recombinant WT or N615S receptors under 1 mM $Mg^{2+}$ conditions

| Figure | Endpoint / dataset | *n* values | Comparison or effect | Test | Statistic (df) | *P* value |
|---|---|---|---|---|---|---|
| Fig. 4h | Relative response, HEK293T cells | Total *n* = 49 cells | Genotype effect | Two-way ANOVA | F(1,43) = 522.222 | <0.001 |
| Fig. 4h | Relative response, HEK293T cells | Total *n* = 49 cells | Drug effect | Two-way ANOVA | F(2,43) = 46.617 | <0.001 |
| Fig. 4h | Relative response, HEK293T cells | Total *n* = 49 cells | Genotype × drug interaction | Two-way ANOVA | F(2,43) = 7.815 | 0.001 |
| Fig. 4h | Relative response, HEK293T cells | Mem: WT *n* = 12, N615S *n* = 6 | WT vs N615S, memantine | Holm–Sidak *post hoc* test | t = 16.557 | <0.001 |
| Fig. 4h | Relative response, HEK293T cells | Ket: WT *n* = 5, N615S *n* = 8 | WT vs N615S, ketamine | Holm–Sidak *post hoc* test | t = 12.555 | <0.001 |
| Fig. 4h | Relative response, HEK293T cells | K2060: WT *n* = 5, N615S *n* = 13 | WT vs N615S, K2060 | Holm–Sidak *post hoc* test | t = 10.300 | <0.001 |
| Fig. 4h | Relative response, HEK293T cells | WT: Mem *n* = 12, Ket *n* = 5 | Memantine vs ketamine, WT | Holm–Sidak *post hoc* test | t = 4.912 | <0.001 |
| Fig. 4h | Relative response, HEK293T cells | WT: Mem *n* = 12, K2060 *n* = 5 | Memantine vs K2060, WT | Holm–Sidak *post hoc* test | t = 9.269 | <0.001 |
| Fig. 4h | Relative response, HEK293T cells | WT: Ket *n* = 5, K2060 *n* = 5 | Ketamine vs K2060, WT | Holm–Sidak *post hoc* test | t = 3.667 | 0.002 |
| Fig. 4h | Relative response, HEK293T cells | N615S: Mem *n* = 6, Ket *n* = 8 | Memantine vs ketamine, N615S | Holm–Sidak *post hoc* test | t = 2.765 | 0.017 |
| Fig. 4h | Relative response, HEK293T cells | N615S: Mem *n* = 6, K2060 *n* = 13 | Memantine vs K2060, N615S | Holm–Sidak *post hoc* test | t = 4.205 | <0.001 |
| Fig. 4h | Relative response, HEK293T cells | N615S: Ket *n* = 8, K2060 *n* = 13 | Ketamine vs K2060, N615S | Holm–Sidak *post hoc* test | t = 1.295 | 0.202 |

Supplementary Table S10
Statistical analysis of OFT (related data are shown in Figure 5)

| Figure | Endpoint/dataset | Comparison or effect | Test | Statistic (df) | *P* value |
|---|---|---|---|---|---|
| Fig. 5c | Distance travelled over time | Genotype effect | Two-way repeated-measures ANOVA | F(1,18)=128.505 | <0.001 |
| Fig. 5c | Distance travelled over time | Time-bin effect | Two-way repeated-measures ANOVA | F(7,126)=0.795 | 0.593 |
| Fig. 5c | Distance travelled over time | Genotype x time-bin interaction | Two-way repeated-measures ANOVA | F(7,126)=5.669 | <0.001 |
| Fig. 5d | Distance travelled in WT after K2060 | Dose effect | Two-way repeated-measures ANOVA | F(4,39)=1.019 | 0.410 |
| Fig. 5d | Distance travelled in WT after K2060 | Time-bin effect | Two-way repeated-measures ANOVA | F(7,273)=48.425 | <0.001 |
| Fig. 5d | Distance travelled in WT after K2060 | Dose x time-bin interaction | Two-way repeated-measures ANOVA | F(28,273)=1.133 | 0.299 |
| Fig. 5f | Distance travelled in N615S after K2060 | Dose effect | Two-way repeated-measures ANOVA | F(3,30)=10.276 | <0.001 |

| | | | | | |
|---|---|---|---|---|---|
| Fig. 5f | Distance travelled in N615S after K2060 | Time-bin effect | Two-way repeated-measures ANOVA | F(7,210)=0.645 | 0.718 |
| Fig. 5f | Distance travelled in N615S after K2060 | Dose x time-bin interaction | Two-way repeated-measures ANOVA | F(21,210)=2.154 | 0.003 |

Supplementary Table S11
Statistical analysis of ASR and PPI data (related data are shown in Figure 5)

| Figure | Endpoint/dataset | Comparison or effect | Test | Statistic (df) | P value |
|---|---|---|---|---|---|
| Fig. 5g | ASR amplitude | Genotype effect | Mann–Whitney rank-sum test | U = 6.000 | <0.001 |
| Fig. 5h | Baseline PPI | Genotype effect | Two-way repeated-measures ANOVA | F(1,28) = 3.311 | 0.080 |
| Fig. 5h | Baseline PPI | Genotype × prepulse intensity interaction | Two-way repeated-measures ANOVA | F(3,84) = 5.282 | 0.002 |
| Fig. 5h | Baseline PPI | WT vs N615S at +16 dB | Holm–Sidak *post hoc* test | t = 3.601 | <0.001 |
| Fig. 5i | PPI after K2060 treatment in WT mice | Treatment effect | Two-way repeated-measures ANOVA | F(1,14) = 1.534 | 0.236 |
| Fig. 5i | PPI after K2060 treatment in WT mice | Treatment × prepulse intensity interaction | Two-way repeated-measures ANOVA | F(3,42) = 1.621 | 0.199 |
| Fig. 5j | PPI after K2060 treatment in N615S mice | Treatment effect | Two-way repeated-measures ANOVA | F(1,12) = 0.041 | 0.843 |
| Fig. 5j | PPI after K2060 treatment in N615S mice | Treatment × prepulse intensity interaction | Two-way repeated-measures ANOVA | F(3,36) = 0.888 | 0.457 |

PPI data were analysed using two-way repeated-measures ANOVA. For baseline PPI, genotype was used as the between-subject factor and prepulse intensity as the repeated factor. For treatment experiments, vehicle and K2060 groups were analysed separately within each genotype, with treatment as the between-subject factor and prepulse intensity as the repeated factor. The main effect of prepulse intensity is not shown because it reflects the expected increase in PPI with increasing prepulse intensity. Holm–Sidak *post hoc* tests were used where appropriate. ASR amplitude was analysed using the Mann–Whitney rank-sum test because normality was not met.

Supplementary Table S12
Statistical analysis of audiogenic seizure test outcomes (related data are shown in Figure 6)

| Figure | Dose | Treatment | Normal response | Abnormal response | Comparison | Test | P value |
|---|---|---|---|---|---|---|---|
| Fig. 6a | 5 mg/kg | Vehicle | 0 | 8 | Reference | Fisher's exact test | — |
| Fig. 6a | 5 mg/kg | K2060 | 7 | 1 | vs dose-matched vehicle | Fisher's exact test | 0.0014 |
| Fig. 6a | 5 mg/kg | Memantine | 7 | 1 | vs dose-matched vehicle | Fisher's exact test | 0.0014 |
| Fig. 6a | 5 mg/kg | Ketamine | 8 | 0 | vs dose-matched vehicle | Fisher's exact test | <0.001 |
| Fig. 6b | 1 mg/kg | Vehicle | 0 | 7 | Reference | Fisher's exact test | — |
| Fig. 6b | 1 mg/kg | K2060 | 5 | 2 | vs dose-matched vehicle | Fisher's exact test | 0.021 |
| Fig. 6b | 1 mg/kg | Memantine | 2 | 5 | vs dose-matched vehicle | Fisher's exact test | 0.462 |
| Fig. 6b | 1 mg/kg | Ketamine | 1 | 7 | vs dose-matched vehicle | Fisher's exact test | 1.000 |

For statistical analysis, behavioural outcomes were dichotomised as normal response versus abnormal audiogenic response. WR, AGS and AGS-RA were pooled as abnormal responses. Each treatment group was compared with the corresponding dose-matched vehicle group using Fisher's exact test.
